# Structure of the human mitochondrial RNA degradosome reveals a distinct mode of helicase–nuclease coupling

**DOI:** 10.64898/2026.09.06.749684

**Authors:** Paula Favoretti Vital do Prado, Saruby Sharma, Arjun Bhatta, Ana Vučković, Maral Parvizian Marani, Andrei Phillip Llarena David, Bernhard Kuhle, Hauke S. Hillen

## Abstract

RNA degradation shapes cellular and organellar gene expression. In human mitochondria, this process is mediated by a dedicated degradosome comprising the helicase SUV3 and the exoribonuclease PNPase, but how these enzymes assemble and cooperate to degrade structured RNA has remained unknown. Here, we report the cryo-EM structure of the complete human mitochondrial RNA degradosome bound to RNA. The structure reveals an asymmetric heteropentamer composed of a SUV3 dimer and the trimeric PNPase. Degradosome assembly is accompanied by conformational rearrangements in PNPase that reshape the RNA-entry channel and generate an asymmetric S1-domain platform for SUV3 binding. This creates a continuous electropositive path from SUV3 to PNPase, suggesting how RNA may be guided during degradation. Together, these findings reveal a mode of helicase–nuclease coupling distinct from other RNA degradation machineries and provide a structural framework for understanding human mitochondrial RNA degradation, its regulation, and association with disease.

## INTRODUCTION

Balancing the rates of RNA synthesis and degradation is one of the universal processes underlying the regulation of cellular gene expression. In human mitochondria, RNAs are transcribed from a dedicated mitochondrial genome (mtDNA) into near-genome length polycistronic transcripts that are subsequently processed to generate mitochondrial tRNAs, rRNAs, and mRNAs ^1–6^. As a result, mt-RNAs are initially produced in stoichiometric amounts. Yet, the steady-state levels of mt-mRNAs vary substantially ^7^, indicating a major role for post-transcriptional mechanisms in the regulation of mitochondrial gene expression. RNA degradation and its interplay with mt-RNA synthesis, maturation, and stabilization has emerged as one of the central processes shaping the mitochondrial transcriptome ^8,9^, from balancing mt-mRNA abundances to eliminating spurious, aberrant, or faulty transcripts that would otherwise interfere with correct mitochondrial function ^10,11^. In addition, mitochondrial RNA degradation may also play a role in immune regulation ^12^.

Throughout all kingdoms of life, RNA degradation is carried out by multi-protein assemblies in which a central ribonuclease is supported by accessory factors that mediate substrate recognition and selectivity and enhance processing efficiency ^13,14^. In archaea and the eukaryotic nucleo-cytoplasmic compartments RNA processing and decay are mediated primarily by the exosome, a conserved multi-protein ribonuclease complex ^15,16^. In bacteria, these activities are organized within the RNA degradosome, in which the exoribonuclease polynucleotide phosphorylase (PNPase) associates with RNase E, the helicase RhlB, and additional cofactors ^17–20^. Among the accessory factors, helicases play a key role by unwinding and thus facilitating the degradation of structured RNA regions ^21^. The minimal realization of this principle is represented by the yeast mitochondrial degradosome (mtEXO), a 1:1 assembly of the nuclease Dss1 and the helicase SUV3 that exhibit strong functional interdependence ^22^.

In human mitochondria, RNA degradation is mediated by a dedicated mitochondrial RNA degradosome, which is comprised of the nuclease PNPase (hPNPase) and the helicase SUV3 (hSUV3). SUV3 is an ATP-dependent RNA/DNA helicase of the SF2 family with 3′→5′ directionality ^23–25^. PNPase, a homolog of bacterial PNPases, employs inorganic phosphate to catalyze the processive 3′→5′ phosphorolysis of RNA, generating nucleoside diphosphates ^26^. Biochemical studies have demonstrated that these enzymes assemble into a heteropentameric complex that degrades double-stranded RNA (dsRNA) substrates bearing single-stranded 3′ overhangs in an ATP-dependent manner ^27^. Depletion of degradosome components leads to the accumulation of mitochondrial dsRNA (mt-dsRNA) ^28,29^. In the case of PNPase depletion, this is accompanied by the release of mt-dsRNA into the cytosol, where it triggers an interferon response ^29^. Loss of mitochondrial degradosome function also results in the buildup of other aberrant RNA species, including nonsense transcripts and R-loops, leading to compromised genome stability and transcriptional dysregulation ^10,11^. Consistent with these findings, mutations in the *PNPT1* and *SUPV3L1* genes that encode PNPase and SUV3, respectively, have been associated with severe human disease ^30–45^.

Despite its fundamental role in organellar gene expression, the structural and molecular basis of RNA degradation in human mitochondria remains unknown. Previous studies have reported structures of human SUV3 in its monomeric form ^23,46^ and, very recently, also in the dimeric state ^47^, which is presumed to represent the physiologically active form. Structures of the trimeric PNPase both in its apo and RNA-bound forms have also been reported ^48–50^. To date, however, no high-resolution structure of the complete degradosome complex is available. It therefore remains unclear how hSUV3 and hPNPase assemble to degrade structured mitochondrial RNAs.

Here, we present the single-particle cryo-EM structure of the complete human mitochondrial degradosome bound to RNA. The structure reveals an asymmetric, heteropentameric complex composed of a dimer of SUV3 and a trimer of PNPase. Their interaction is facilitated by conformational rearrangements in PNPase, which create a binding platform for SUV3 and produce a continuous electropositive groove from the helicase to the nuclease, suggesting a pathway for RNA during processive degradation. Comparison to other known RNA degradation machineries shows that the human mitochondria degradosome employs an architecturally unique mode of helicase-nuclease coupling. Taken together, these findings provide the first structural snapshots of the complete human mitochondrial RNA degradation machinery and establish a framework for future mechanistic studies of mitochondrial RNA decay and its links to disease.

## RESULTS

### Cryo-EM structure of the human mitochondrial degradosome

To investigate the molecular basis of mitochondrial RNA degradation, we recombinantly expressed and purified human SUV3 and PNPase (Extended Data Figure 1a). The degradosome was reconstituted by incubating both components together and complex formation was verified by analytical size-exclusion chromatography (Extended Data Figure 1b). To confirm the biochemical activity of the complex, we incubated PNPase and SUV3 with a fluorescently labeled RNA comprising a double-stranded segment and a single-stranded 3’ overhang in the presence of magnesium and phosphate ^50–52^ (Extended Data Figure 1c, Extended Data Table 1). Consistent with previous reports ^27^, PNPase alone efficiently degraded a single-stranded RNA probe but only partially degraded the substrate with a double-stranded segment (Extended Data Figure 1d). By contrast, incubation of this substrate with both PNPase and SUV3 in the presence of ATP resulted in near-complete degradation of the dsRNA substrate, indicating that SUV3 facilitates degradation of the double-stranded RNA (Extended Data Figure 1d) ^27,46^. Together, these results confirm that the reconstituted human mitochondrial degradosome is functional.

To stabilize the complex for structural analysis, we designed an RNA substrate comprising a 60-nt single-stranded RNA annealed to a 10-nt complementary oligonucleotide, generating a double-stranded region with a 50-nt single-stranded 3′ overhang (Extended data Table 1 – dsRNA_02_PT). The longer RNA strand additionally contained five phosphorothioate linkages, which confer resistance to nucleolytic degradation (Extended Data Figure 1e – dsRNA_02_PT) ^50–52^. Biochemical assays confirmed that the SUV3–PNPase complex failed to completely degrade the modified substrate, even in the presence of ATP (Extended Data Figure 1f).

SUV3 and PNPase were incubated with this substrate in the absence of Mg^2+^ and ATP and the resulting complex was used for single particle cryo-EM analysis. After image processing, this led to a reconstruction of the human mitochondrial degradosome at a global resolution of 3.3 Å (Extended Data Figure 2, Methods). In the consensus reconstruction, the density for SUV3 was less well-defined than for PNPase, indicating conformational flexibility relative to PNPase. To overcome this, we used RELION multibody refinement ^53^, which yielded improved maps with local resolutions of 3.6 Å for SUV3 and 3.2 Å for PNPase. Using previously reported structures of SUV3 ^46,54^ and PNPase ^49,50^ as well as AlphaFold3 ^55^ models, we were able to build and refine an atomic model of the human mitochondrial degradosome with excellent stereochemistry (Extended Data Table 2).

The resulting structure reveals the architecture of the human mitochondrial degradosome. It is composed of a trimer of PNPase (PNPase a-c) and a dimer of SUV3 (SUV3 a-b), which together form a heteropentameric complex, as previously suggested ^27^ (Figure 1). Consistent with earlier SAXS studies ^46^, the complex adopts a dumbbell-shaped architecture, with the SUV3 dimer positioned on top of the trimeric PNPase base. The interaction between the two subcomplexes is mediated by the S1 domains of PNPase, which form a chalice-like docking site that interacts with both SUV3 subunits (Extended Data Figure 3a-b). Additional density corresponding to RNA is visible at multiple locations within the complex, including the active site of SUV3-a, near the exit channel of SUV3-b extending toward PNPase S1-a, and at the entrance to the PNPase catalytic chamber (Extended Data Figure 3c-d).

**Figure 1.**
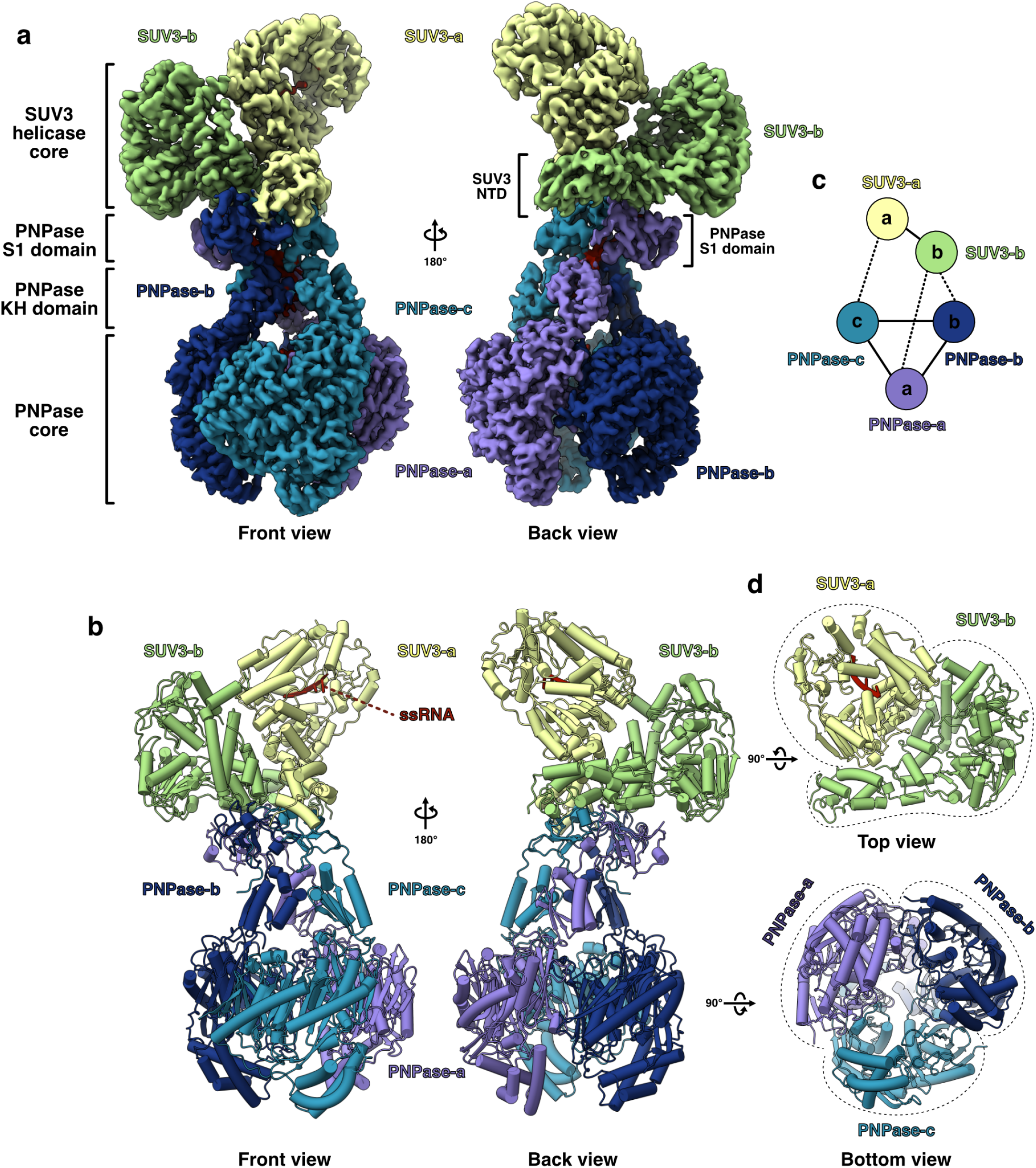
Structure of the human mitochondrial RNA degradosome. **a)** Composite cryo-EM map (Map D) of the degradosome colored by subunit. The two SUV3 subunits are shown in shades of green; PNPase subunits are shown in shades of blue and purple; RNA density is shown in red. The helicase and exonuclease cores are indicated. **b)** Cartoon representation of the degradosome, colored as in (A). α-Helices are depicted as cylinders. **c)** Chain-contacts diagram. Nodes represent individual chains, and lines indicate inter-chain interfaces based on buried surface area. Dashed lines represent smaller interfaces. **d)** Top and bottom views of the complex, highlighting the top of the SUV3 dimer and the bottom of the PNPase trimer architecture.

### SUV3 forms an asymmetric dimer

The structure of the degradosome captures SUV3 in its active, dimeric state (Figure 2). SUV3 consists of an N-terminal domain (NTD) followed by a conserved catalytic core composed of two RecA-like domains (RecA1 and RecA2) arranged in tandem and a C-terminal domain (CTD) with a C-terminal tail (CTT, residues 723–786) (Figure 2a). The C-terminal domain folds opposite the RecA domains, forming a ring-like structure through which the RNA strand is threaded ^23,46^. The two SUV3 subunits form an asymmetrical homodimer in which both adopt an identical overall fold and spatial arrangement (r.m.s.d. = 0.601 across 591 pruned Cα-atom pairs) (Extended Data Figure 4a). The two monomers interact in a concave-convex fashion, with the convex surface of SUV3-a accommodated within the concave surface of SUV3-b (Figure 2c-d). Dimerization is mediated by three main interfaces (Figure 2c). The first is formed between the NTD and CTD of SUV3-a and the CTD of SUV3-b. The second is mediated by the SUV3-a CTD and the SUV3-b RecA2 domain. The third involves the SUV3-a RecA1 domain and the NTD of SUV3-b. Comparison to the structure of the free SUV3 dimer ^47^, which was reported during the preparation of this manuscript, shows that while the overall architecture is similar, the NTDs of both SUV3 copies adopt distinct orientations in the degradosome (Extended Data Figure 4b). Previous studies have reported that the CTT of SUV3 is required for dimerization ^46,47^. Surprisingly, this region is not resolved in either the free SUV3 dimer ^47^ or the degradosome structure, suggesting that it remains disordered or highly flexible even upon PNPase binding.

**Figure 2.**
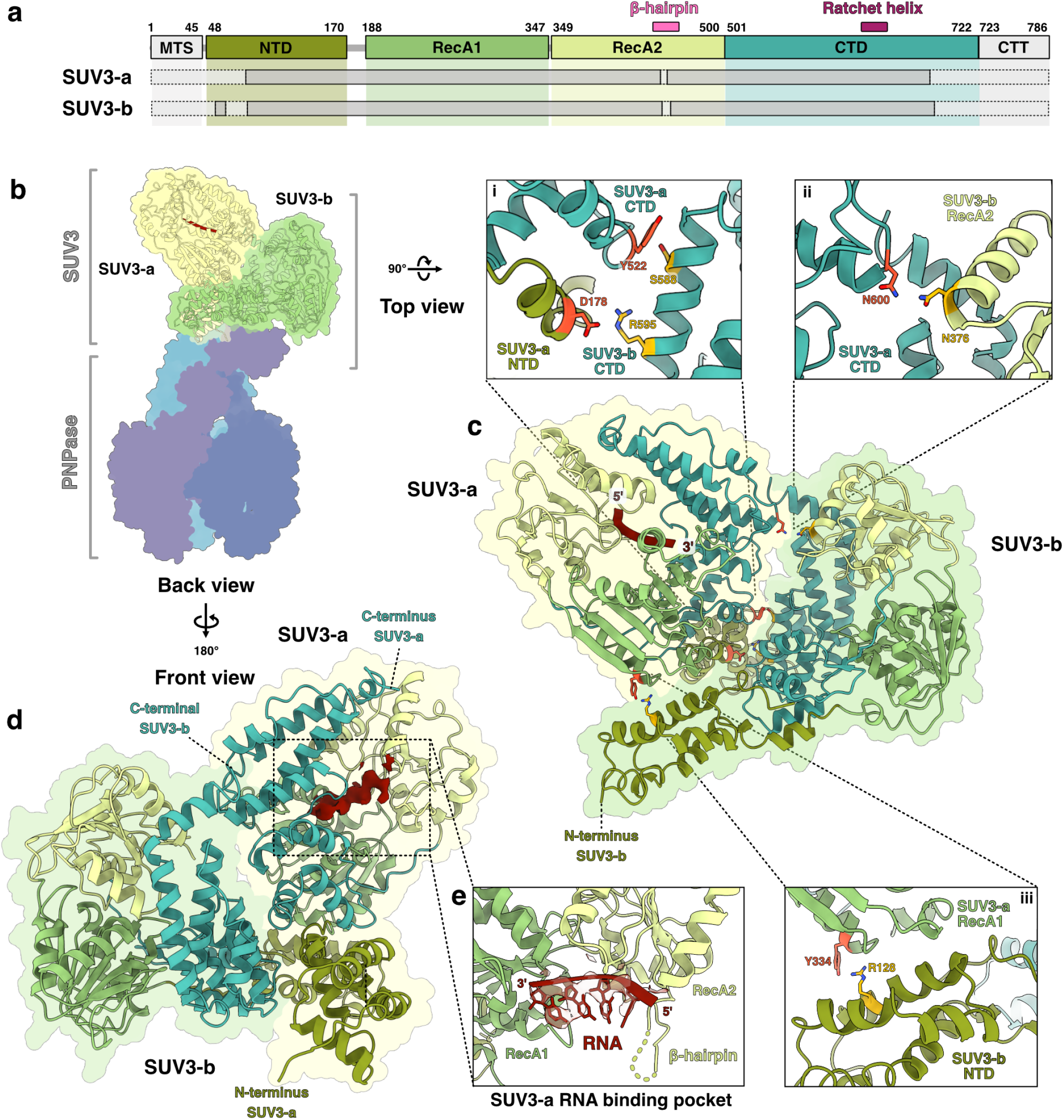
Structure of the SUV3 dimer. **a)** Domain representation of SUV3. For each subunit, modeled regions are indicated by blocks with solid black lines; segments not modeled in the structure are shown as transparent blocks with dashed lines. Conserved elements of the translocation module are annotated on top. **b)** Back view of the degradosome complex with the SUV3 dimer indicated and shown as cartoon. PNPase subunits are shown as transparent surfaces. **c)** Top view of the degradosome with PNPase subunits omitted for clarity. The SUV3 dimer is shown as cartoon representations, colored by domain as in (A). Insets (i–iii) highlight detailed views of the three principal regions mediating dimer-stabilizing interactions, with interacting domains and residues indicated (residues within 3 Å, as identified using ChimeraX). **d)** Enlarged front view of the complex; PNPase subunits are omitted for clarity. The SUV3 dimer is shown as cartoons colored by domain; RNA density is shown in red. **e)** Poly(U) ssRNA in SUV3-a. The disordered β-hairpin is indicated by dashed lines, the CTD is omitted for clarity.

The active site of SUV3-a contains additional density consistent with four nucleotides of RNA (Figure 2d-e). Structural comparison shows that the RNA occupies the same position as in the previously reported RNA-bound SUV3 structures (Extended Data Figure 4b-c) ^23,47^. Because the quality of the map in this region is insufficient to distinguish individual bases, we modeled the RNA as poly(U). The 3′ end of the modeled RNA points toward the dimer interface, facing SUV3-b (Figure 2c). However, no further density corresponding to RNA is visible within the SUV3 dimer and the active site of SUV3-b appears unoccupied (Extended Data Figure 4d). These observations are consistent with the recent structure of the free SUV3 dimer bound to RNA, in which similarly only one subunit engages RNA ^47^.

To obtain further insight into the potential mechanism of SUV3, we carried out a structural similarity search using Foldseek ^56^. Among the highest scoring hits was Hel308 (E-value 7.33e-13), an archaeal SF2 helicase whose structure has been determined in complex with a DNA duplex–single-strand junction, providing detailed insight into substrate engagement during unwinding ^57^. In the Hel308–DNA complex, the RecA-CTD ring accommodates the 3′ end of the single-stranded unwound product in an ATP-independent manner (Extended Data Figure 4e). The nucleic acid backbone engages with the RecA domains and is stabilized by base-stacking interactions with a central helix, which acts as a ratchet that promotes directional translocation along the nucleic acid (ratchet helix) ^57^. Together, the RecA domains and the ratchet helix constitute the “translocation module”, which binds and moves along the loaded strand of the RNA (Extended Data Figure 4e). The entry to the translocation module is gated by a β-hairpin in RecA2 that acts as a wedge to split the DNA duplex (Extended Data Figure 4e-f) ^57^. Structural comparisons suggest that key features of this architecture are conserved in SUV3 (Extended Data Figure 4f). First, SUV3 contains a structurally equivalent element to the β-hairpin (residues 441– 461) (Extended Data Figure 4f). This loop is poorly defined in the cryo-EM density, suggesting conformational flexibility. Second, the helix spanning SUV3 residues 441–461 may be structurally and functionally homologous to the ratchet helix (Extended Data Figure 4f). Remarkably, while Hel308 acts as a monomer, the two translocation modules of the SUV3 dimer are arranged in tandem and share the same directionality, raising the possibility that they form an extended protein–nucleic acid interaction pathway (Extended Data Figure 4g).

Taken together, the degradosome structure demonstrates that SUV3 adopts a similar dimeric assembly in complex with PNPase as in its free form, with RNA binding to only one of its subunits. Moreover, comparison with Hel308 indicates that key structural features required for nucleic acid unwinding and translocation are conserved, suggesting that SUV3 may operate through a similar mechanism.

### Structural rearrangements prime PNPase for processive RNA degradation

The degradosome structure shows that PNPase adopts a unique conformation in this complex. PNPase assembles into a homotrimeric complex, forming a toroidal (doughnut-shaped) structure in which six PH–like domains delineate a central channel ^48–50^. Within each monomer, two RNase PH domains (RNase PH1 and RNase PH2) are connected by an all-α-helical domain and form the exonuclease core (Figure 3a). RNase PH2 is followed by the S1 and KH domains, two RNA-binding modules that together form a “cap domain” that crowns the nuclease core and forms an RNA entry channel (Figure 3a-c) ^49^.

Comparisons to previous structures of PNPase in the apo and RNA-bound states show that key elements of PNPase are reorganized within the degradosome. First, the S1 domains of PNPase adopt a unique, asymmetric conformation (Figure 3d-e).

**Figure 3.**
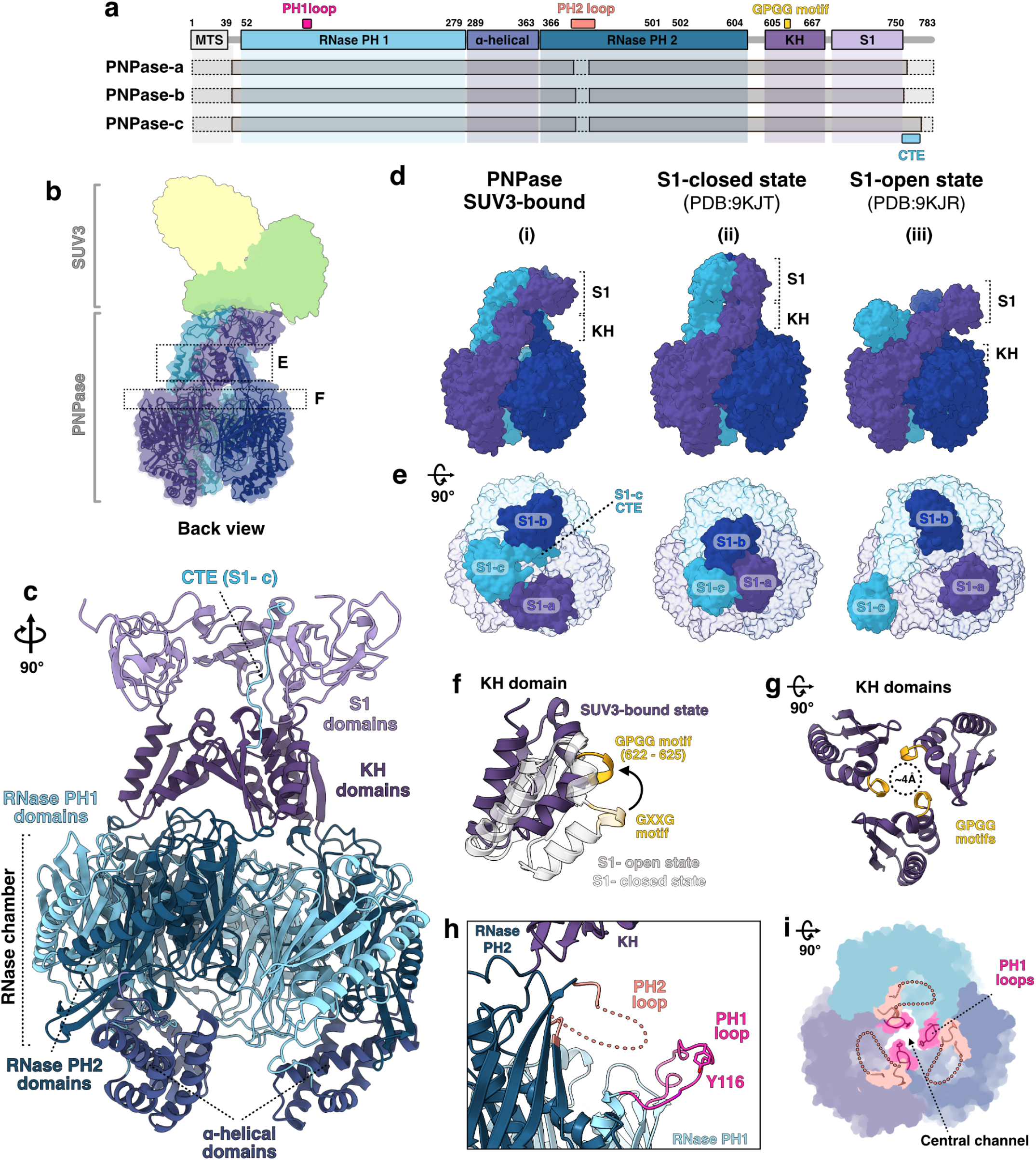
PNPase structure within the degradosome. **a)** Domain representation of PNPase. For each subunit, modeled regions are indicated by solid black lines; segments not modeled in the structure are shown as dashed lines. The PH1 loop (and the YLRR motif), PH2 loop and GPGG motif are indicated above. **b)** Front view of the degradosome with PNPase shown as a cartoon; SUV3 subunits are shown as transparent surfaces. Dashes boxes indicate cross sections of the regions enlarged in F and G. **c)** PNPase colored by domains as in(A). The view is rotated relative to panel B. **d)** Side view of PNPase S1-domain flexibility shown side-by-side: (i) SUV3-bound PNPase (this work), SUV3 is omitted for clarity. (ii) PNPase S1-closed state (PDB: 9KJT ^49^), (iii) PNPase S1-open state (PDB: 9KJR ^49^). Structures are shown as surfaces; subunits are colored by chain as in Figure 1. **e)** Top view of PNPase S1-domain flexibility shown side-by-side: (i) SUV3-bound PNPase (this work), SUV3 is omitted for clarity. (ii) PNPase S1-closed state (PDB: 9KJT ^49^), (iii) PNPase S1-open state (PDB: 9KJR ^49^). Structures are shown as surfaces; the view is rotated relative to panel B; subunits are colored by chain as in Figure 1. **f)** Representative view of KH domain repositioning. The PNPase catalytic core was aligned in both open and closed states ^49^, with the isolated KH-c domain shown as an example of the conformational changes that reposition the loop containing the GXXG motif. **g)** Rearrangements of the KH domains in the degradosome constrict the central channel. The view is rotated relative to panel B. **h)** Relative positions of the PH2 loop and PH1 loop in one of the PNPase protomers of the degradosome. The PH2 loop is represented by dashed lines. **i)** Geometry of the central channel in degradosome PNPase. The disordered PH2 loop is represented by dashed lines. The view is rotated relative to panel B.

Previous structures have shown that the S1 domains can undergo substantial movements relative to the exonuclease core, transitioning from a closed state, in which the three S1 domains symmetrically interact with one another, to an open state, in which the S1 domains are positioned farther apart from each other and closer to the RNase PH2 domains of adjacent subunits (Figure 3d-e) ^49^. In the degradosome, the S1 domains adopt an asymmetric arrangement not previously observed, in which S1-c and S1-b establish extensive interactions with each other while S1-a is disengaged (Figure 3d-e).

Second, the KH domains adopt a unique orientation with respect to the exonuclease core (Figure 3f). In all previously reported structures, this domain adopts a bent-inward orientation and engages with a loop protruding from the RNase PH2 domain (PH2 loop; residues 402–423) ^48–50^. By contrast, in the degradosome, none of the three KH domains establishes contacts with the RNase core. Instead, all three adopt outward-bent conformations and are slightly rotated (Figure 3f), thereby altering the geometry of the RNA-entry channel (Figure 3g, Extended Data Figure 5b). This positions a glycine-rich loop upward and inward toward the central channel, exposing a conserved GXXG motif (GPGG in human PNPase, residues 622-625) implicated in RNA interactions in both bacterial and human PNPase ^49,58^. As a result, the GXXG motif is directed toward the trimer core, resulting in a constricted channel aperture (Figure 3g). This is reminiscent of bacterial PNPase ^59^, where the KH domains undergo conformational changes upon substrate binding and adopt a constricted conformation that clamps the RNA (Extended Data Figure 5a).

Finally, the PH1 and PH2 loops, two mobile elements that line the RNA-entry channel leading to the RNase PH chamber, adopt distinct conformations (Figure 3h-i, Extended Data Figure 5c-d). The PH2 loop, located immediately below the base of the KH domain, functions as a gate to the degradation chamber. In apo PNPase, this loop adopts two major conformational states: an open state, in which it points outward toward the neighboring PNPase subunit, and a closed state, in which it is oriented toward the center of the pore (Extended Data Figure 5d-e) ^49^. These conformations expose or occlude the central channel of the RNase PH chamber, respectively (Extended Data Figure 5d-e). Recent structures of PNPase bound to ssRNA ^50^ have shown that the PH2 loop can further undergo dynamic transitions among multiple conformations within its closed state (Extended Data Figure 5f). This permits the neighboring PH1 loops (residues 104–124) to adopt an asymmetric conformation, with one of them extending into the active site of an adjacent subunit forming contacts between Y116 and R119 and active site residues L443 and N444. This was suggested to stabilize the active site and restrict activity to one of the three active sites per catalytic cycle ^50^. In the degradosome, the PH2 loop shows no well-defined density, suggesting conformational flexibility (Figure 3h-i). Furthermore, the PH1 loop does not engage in interactions with active site residues. Instead, Y116 faces the central pore, occupying a position equivalent to that of F77 in RNA-bound bacterial PNPases (Extended Data Figure 5g-h). The latter has been shown to contribute to proper substrate positioning within the central channel and to facilitate RNA guidance into the catalytic cavity through base-stacking interactions ^58–60^. Thus, Y116 of human PNPase may be functionally equivalent to F77 in bacterial PNPase. Consistent with this, we observe density that we attribute to RNA within the pore (Extended Data Figure 5g).

In summary, PNPase adopts a distinct conformation within the degradosome, in which rearrangements of the S1, KH, and gating loops reshape the entry channel to support RNA binding.

### Interaction between SUV3 and PNPase

The structural rearrangements in PNPase also provide the basis for its interaction with SUV3. The asymmetric arrangement of S1 domains gives rise to a chalice-shaped binding platform which precisely accommodates the SUV3 dimer (Figure 4a-c). The main interaction interface is formed by S1-b and S1-c, which interact with each other extensively. The C-terminal extension (CTE) of PNPase-c, which is largely disordered in the other two PNPase subunits, adopts a defined conformation and runs along S1-b towards the KH domain of PNPase-b and contacts both SUV3 monomers (Figure 4d). Both S1-b and S1-c establish extensive interactions with the N-terminal domain (NTD) of SUV3-a (Figure 4b-d). Weak density indicates that the CTE of S1-b may also contact the SUV3-a NTD, but we refrained from modeling this. In addition, S1-a interacts with the CTD of SUV3-b (Figure 4c-e). The S1 domains thus interact with both subunits of the SUV3 dimer, explaining previous biochemical data showing that the S1 domains are essential for the interaction between SUV3 and PNPase ^49^ and that SUV3 dimerization is required for degradosome complex formation ^46^.

**Figure 4.**
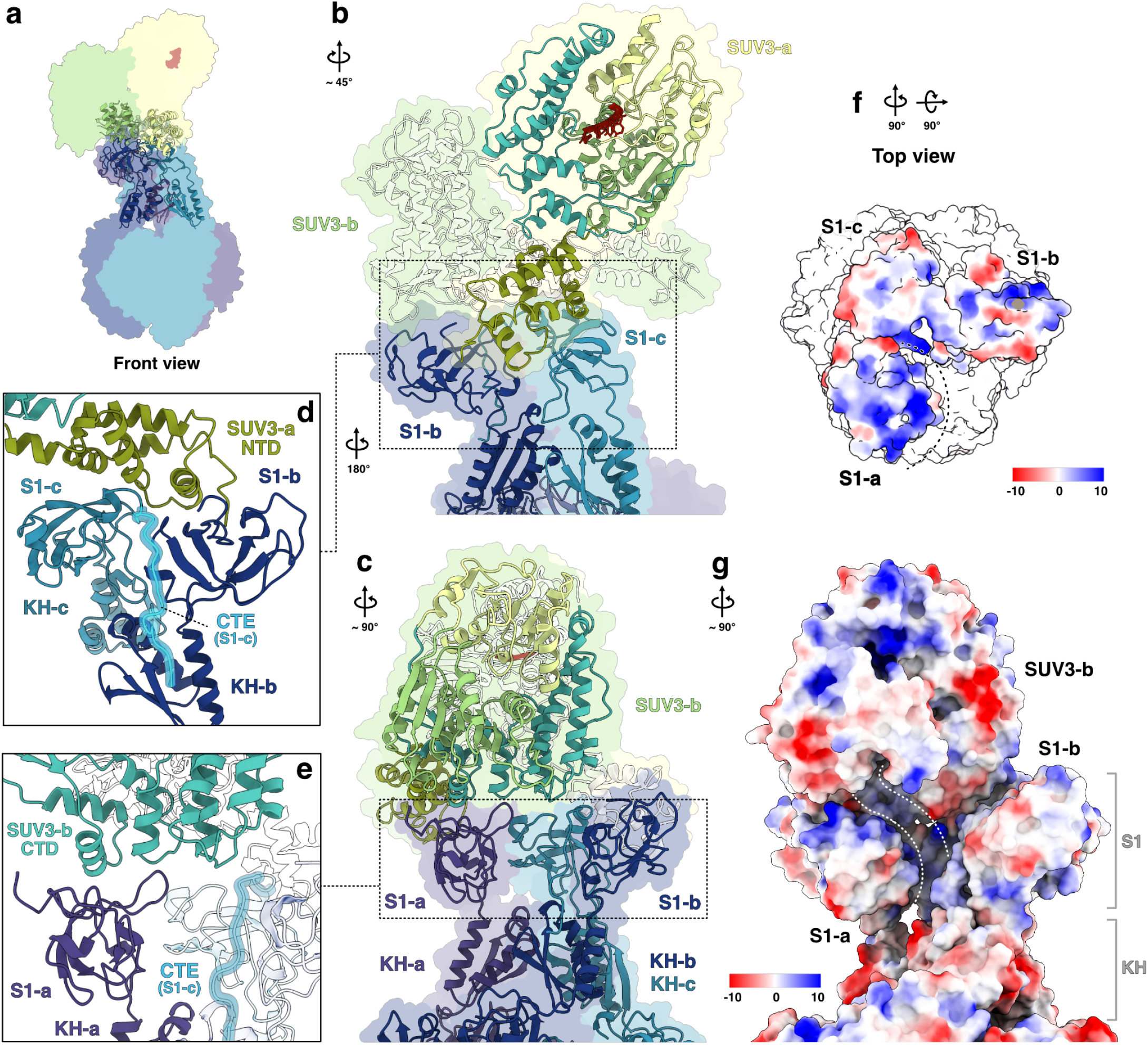
PNPase–SUV3 interactions. **a)** Front view of the degradosome. SUV3 and PNPase domains involved in intermolecular interactions are shown in cartoon presentation. **b)** Overview of the interaction between the SUV3-a NTD and the PNPase S1-b and S1-c domains. For clarity, the SUV3-b subunit is shown as a transparent cartoon. The view is rotated relative to panel A. SUV3-a is colored by domains as in Figure 2. **c)** Overview of the interactions between the SUV3-b subunit and the PNPase S1-a domain. The view is rotated relative to panel A. SUV3-b is colored by domains as in Figure 2. **d)** Close-up of the interactions between the SUV3-a NTD and the PNPase S1-b and S1-c domains. The extended tail of the S1-c domain is highlighted. View is rotated relative to panel B. **e)** Detailed view of the interactions between the SUV3-b CTD and the PNPase S1-a domain. **f)** Top view of the PNPase trimer (SUV3 dimer omitted), shown as surfaces and colored according to Coulombic electrostatic potential (positive surfaces in blue, negative in red). The PNPase core is displayed as a white surface for clarity. **g)** Side view of the degradosome highlighting the cleft generated by the arrangement of the SUV3 dimer with the PNPase trimer. The cleft is delineated with dashed white lines. Surfaces are colored according to Coulombic electrostatic potential (positive surfaces in blue, negative in red). View is rotated relative to A.

The structure also rationalizes a known disease mutation. Mapping of the disease-associated mutation D713Y in PNPase ^29^, which was previously shown to impair binding to SUV3 ^49^, shows that this residue is located directly at the interface between S1-c and SUV3-b (Extended Data Figure 6a-b), suggesting that its mutation may destabilize the interaction.

In addition, the interface between SUV3 and PNPase also suggests a potential path for the RNA. The unique arrangement of the S1 domains positions a positively charged surface patch of S1-a inward toward the center of the trimer (Figure 4f). This creates a groove that contributes to a continuous electropositive surface extending from SUV3-b toward the PNPase RNA entry channel (Figure 4g). Within this groove, we observe additional density that we attribute to RNA (Extended Data Figure 6c-d). However, the quality of the cryo-EM maps in this region does not allow atomic modeling of the RNA.

Taken together, the structure shows that degradosome assembly is facilitated by conformational rearrangements of PNPase, which enable extensive interactions between the PNPase S1 domains and both SUV3 monomers and create an electropositive groove that may guide RNA towards the PNPase degradation chamber.

### A unique mode of helicase-nuclease coupling

The structure reveals that the mitochondrial degradosome employs a unique mode of helicase-nuclease coupling. Structural comparisons to the yeast mitochondrial degradosome and the human exosome show that, although nuclease–helicase coupling is a conserved feature of these machineries, their compositions and architectures vary dramatically (Figure 5a-c). First, both mtEXO and the exosome utilize a monomeric helicase, while the human mitochondrial degradosome contains the dimeric human SUV3. In yeast, the mtEXO complex is a heterodimer composed of a single copy of the SUV3 helicase bound directly to the exonuclease DSS1 ^22^, which is not homologous to PNPase (Figure 5b). The human nuclear RNA exosome consists of a pseudo-hexameric ring of six RNase PH-like proteins capped by three S1/KH domain proteins, which form a catalytically inactive core that resembles the architecture of PNPase (Figure 5c) ^61^. The ribonucleolytic activity is provided by DIS3 (Rrp44), which is positioned at the bottom of the RNase PH-like ring ^61^. The monomeric RNA helicase MTR4 associates with the top of the core through interactions with the S1 domain protein Rrp4 (Figure 5c) ^61^. Second, the interaction interfaces between helicases and nucleases are structurally not conserved in the three complexes. In both mtEXO and the exosome, the interactions between the helicases and DSS1 or MTR4, respectively, differ from those between SUV3 and PNPase in the human mitochondrial degradosome and lead to a different spatial arrangement of the enzymes. Third, the mechanism of RNA handover from helicase to nuclease is different. In mtEXO, the two active sites are directly adjacent and aligned, suggesting a direct handover ^22^. In the exosome, single-stranded RNA is threaded sequentially from MTR4 through the central channel of the core and into the exoribonuclease DIS3 at the opposing side, where degradation occurs ^61^. By contrast, neither of the SUV3 helicase cores in the human degradosome is directly aligned with the central channel of PNPase. Thus, the human mitochondrial degradosome displays a unique architecture and mode of helicase-nuclease coupling.

**Figure 5.**
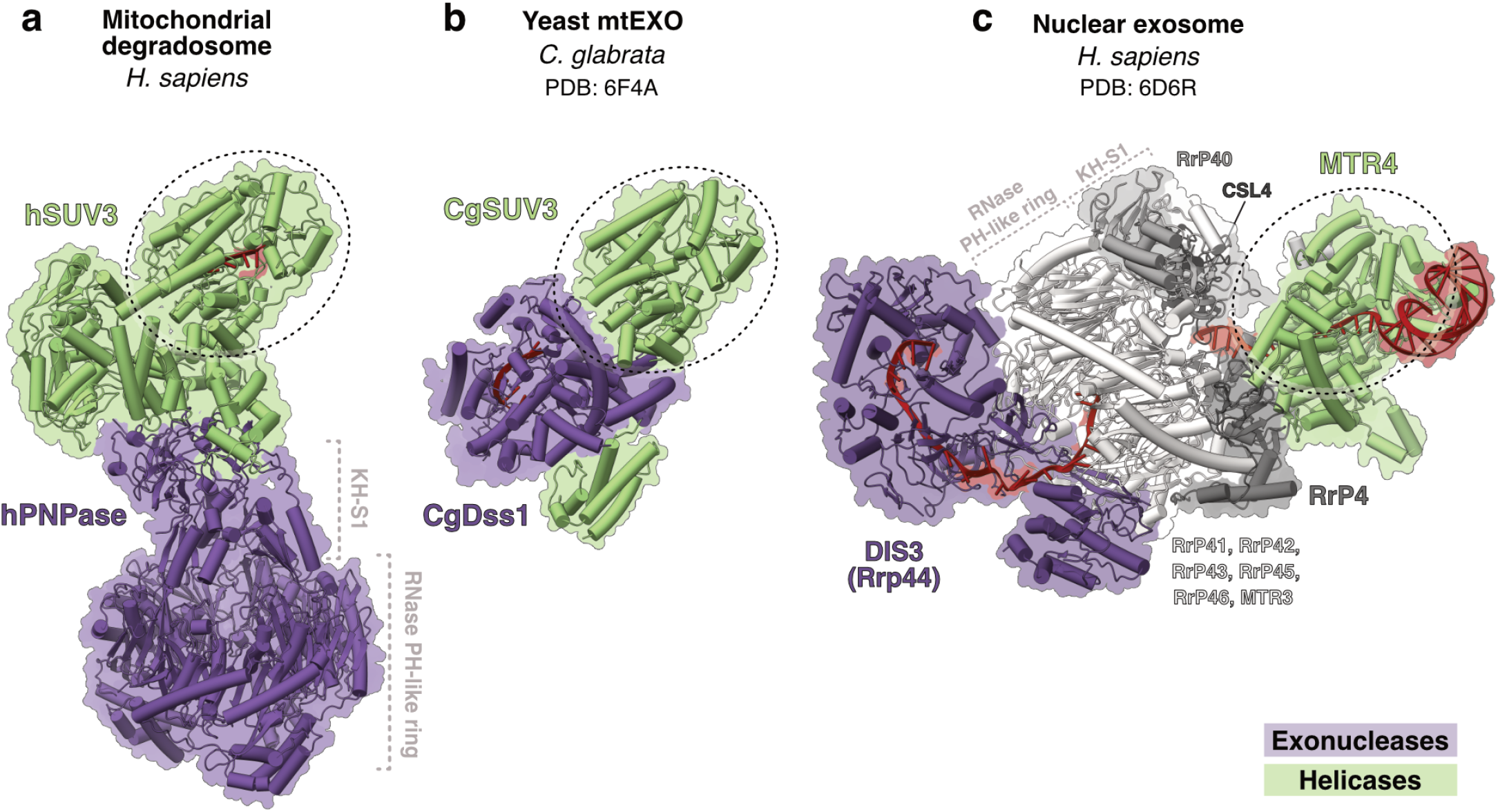
Helicase–nuclease coupling across RNA degradation machineries. Structure of the human mitochondrial degradosome **(a)** shown alongside the yeast mitochondrial degradosome (PDB: 6F4A ^22^) **(b)** and the human nuclear exosome (PDB: 6D6R ^61^) **(c)**. Structures were aligned to SUV3-a and are shown side by side, maintaining the relative orientation established by the alignment, as indicated by the dashed lines. Helicases are colored green, nucleases are colored purple, and additional protein factors are shown in shades of gray. RNA is colored red.

### The path of RNA within the degradosome

The structure of the degradosome also reveals how RNA may be guided from the helicase to the nuclease during processive degradation. While we observe density for RNA in both SUV3 and PNPase, the density is not continuous throughout the complex and in most regions does not allow unambiguous atomic modelling (Figure 6a). Nonetheless, its trajectory suggests potential paths for RNA through the complex.

**Figure 6.**
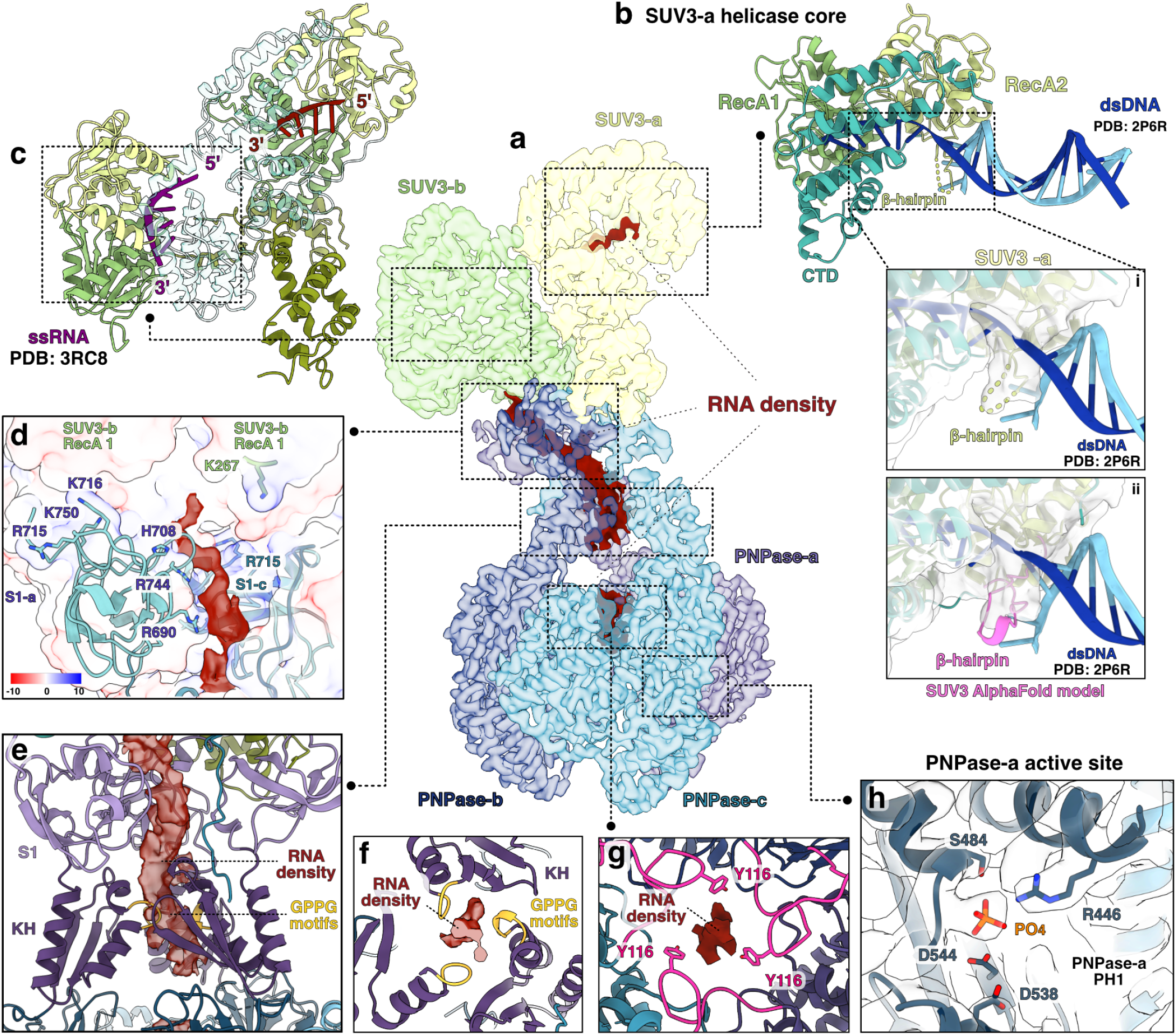
RNA densities throughout the degradosome complex. **a)** Front view of the cryo-EM map of the complex. Protein densities are colored by chain as in Figure 1 and shown with transparency. Additional densities attributed to RNA bound to the complex are shown in solid red. Dashed boxes indicate regions enlarged in subsequent panels. **b)** Model of SUV3 bound with double-stranded DNA (dsDNA). The nucleic acid is positioned by structural superposition with the archaeal helicase Hel308 (PDB: 2P6R ^57^). The unwound DNA strand is shown in dark blue, the complementary strand in light blue. (i) Cryo-EM density around the β-hairpin loop region in SUV3-a. (ii) AlphaFold model of the human SUV3 protomer highlighting the β-hairpin region. **c)** Model of SUV3-b bound to RNA. RNA (shown in purple) was modeled by structural superposition with the human SUV3 monomer bound to short RNA (PDB: 3RC8 ^23^). **d)** Detailed view of the SUV3-b/PNPase S1-a interface. Protein chains are depicted as surfaces colored according to Coulombic electrostatic potential (positive in blue, negative in red). Charged residues in this region are indicated. RNA density is shown in red. **e)** Detailed view of the trace of RNA density spanning from the S1-a domain across the KH domain. The GPGG loop is highlighted in yellow. **f)** Cross-section of the PNPase core at the level of the KH domains, showing RNA density within the central channel surrounded by the GPGG domains. **g)** Cross-section of the PNPase core at the level of the PH1 loop showing RNA density within the central channel, accommodated between the three YLRR loops. Residue Y116 is indicated. **h)** Detailed view of one of the active site of PNPase-a, highlighting the catalytic residues. A phosphate ion was positioned within a globular extra density based on comparison with the apo structure of *C. crescentus* PNPase (PDB: 4AIM ^59^).

During degradation of structured RNA substrates, the RNA is expected to first engage with SUV3, which can translocate along ssRNA and unwind structured regions. Although we do not observe density for a double-stranded RNA segment near SUV3, superposition of SUV3-a with the DNA duplex–bound Hel308 structure demonstrates that the double-stranded nucleic acid could be accommodated without significant steric clashes (Figure 6b) and that the single-stranded 3’ overhang aligns precisely with the RNA observed in the SUV3-a active site (Extended Data Figure 6e). Furthermore, superposition of an AlphaFold3 model of SUV3 that includes the β-hairpin responsible for strand separation in Hel308 positions this feature between the strands at the junction (Figure 6b), consistent with a role in duplex unwinding as proposed for Hel308 ^57^. By contrast, when the Hel308–DNA structure is aligned to SUV3-b, the duplex cannot be accommodated without severe steric clashes (Extended Data Figure 6f). In addition, the β-hairpin of SUV3-b is positioned toward the convex surface of SUV3-a such that one of its β-strands is partially hidden and inaccessible for strand separation. This suggests that SUV3-a may bind an ssRNA-dsRNA junction in a similar fashion as previously observed for Hel308, while SUV3-b likely cannot engage directly with an ss-dsRNA junction.

Surprisingly, we do not observe any RNA density within SUV3-b. However, modelling of an ssRNA into the SUV3-b active site positions the 5′ end facing toward the RNA bound by SUV3-a and the 3′ end towards the positively charged cleft formed by the S1 domains at the interface with PNPase (Figure 6c). This suggests that the ssRNA could potentially be channeled from SUV3-a to SUV3-b during translocation before being directed toward PNPase, consistent with the 3’ to 5’ polarity of RNA degradation by PNPase.

At the SUV3-PNPase interface, we observe RNA density near the interaction region between the NTD–RecA2 ring of SUV3-b and the PNPase S1 domains (Figure 6a and d). The density traces along a positively charged surface of S1-a toward the central channel (Figure 6d) and then extends toward the KH domain, where it passes through the constricted central channel between the GPGG motifs (Figure 6 e-f). This trajectory is consistent with that observed for bacterial PNPase structures bound to RNA, in which the RNA enters the complex from the top with its backbone positioned adjacent to the GXXG loop of the KH domain (Extended Data Figure 5b) ^58,59^, and with previous studies that showed that mutation of the first glycine residue in the GPGG motif of human PNPase (G622D) abolishes RNA binding ^48^. The RNA density then extends to the base of the KH domain and toward the entrance of the PNPase PH1-PH2 catalytic core, where it becomes weak in the region of the disordered PH2 loop, indicating structural flexibility. The density then reappears at the level of the PH1 loop, next to the conserved Y116, supporting an analogous role of this residue in substrate guidance to F77 in bacterial PNPases (Figure 6g) ^58–60^. Beyond this point, the RNA density becomes scattered, with weak density in each of the PNPase active sites, suggesting partial occupancy. However, all three active sites show a clear density consistent with a bound phosphate ion (Figure 6H, Extended Data Figure 6g).

Based on these observations, we propose two potential models of how the RNA may be channelled through the degradosome during processive degradation (Figure 7). In the first model, the substrate RNA is unwound by SUV3 and directly handed over to PNPase for degradation (Figure 7a). The ssRNA-dsRNA substrate is bound by SUV3-a and the unwound ssRNA is channelled through SUV3-b. From there, it runs along the positively charged surface formed by the S1 domains and into the catalytic chamber of PNPase. In the second model (Figure 7b), the RNA is not channelled directly from SUV3 to PNPase and the two enzymes work independently either on the same or different substrate molecules. In this case, SUV3-a may be sufficient for unwinding of ssRNA-dsRNA junctions, and SUV3-b may not directly interact with the RNA. The unwound ssRNA can then re-enter the complex at the junction between SUV3-b and the PNPase S1 domains, which is partially solvent exposed.

**Figure 7.**
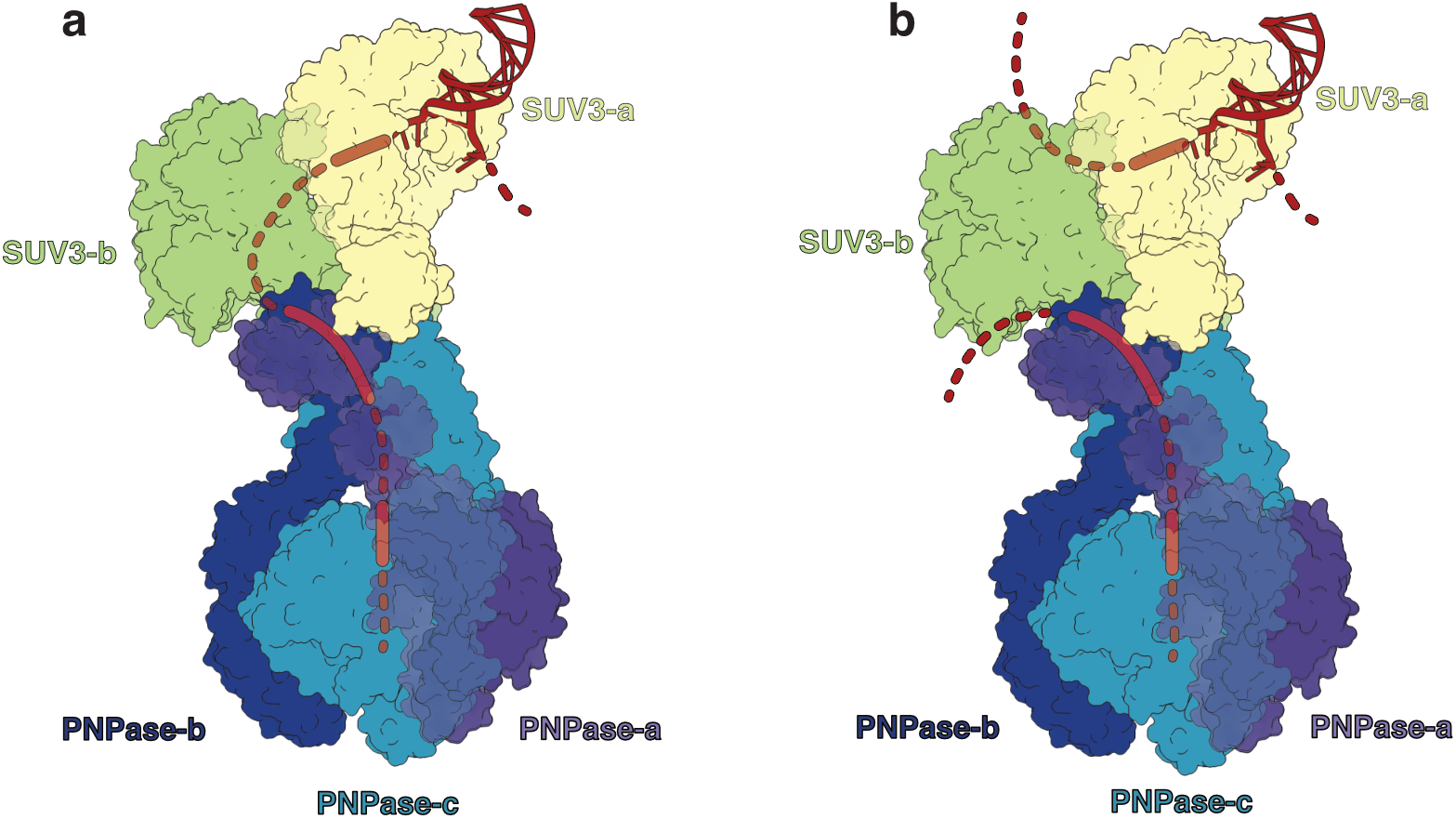
Proposed models for RNA degradation by the degradosome. **a)** Direct channeling model in which ssRNA generated by SUV3-a is transferred through SUV3-b to the PNPase catalytic chamber via the positively charged S1 surface. **b)** Independent activity model in which SUV3 unwinds RNA without direct channeling to PNPase, allowing the released ssRNA to re-enter the complex through the partially solvent-exposed interface between SUV3-b and the PNPase S1 domains.

Taken together, the structure reveals how RNA interacts with SUV3 and PNPase in the degradosome and suggests potential paths for the substrate through the complex.

## DISCUSSION

In this study, we present the structure of the complete human mitochondrial degradosome, which reveals how PNPase and SUV3 interact to mediate RNA degradation. The structure shows how SUV3 dimerizes to adopt its active form, mediated by the NTD, CTD, and RecA domain. Surprisingly, we do not observe any density for the SUV3 CTT, although its deletion has been shown to disrupt dimerization and, consequently, interaction with PNPase ^46^. The CTT may thus facilitate dimer formation by promoting subunit proximity through transient interactions which are not visible in averaging methods such as cryo-EM. Both SUV3 subunits interact directly with PNPase, explaining why its dimerization is essential for degradosome formation ^46^. Structural comparisons suggest that key elements required for dsRNA binding are conserved in SUV3, and that ss-dsRNA junctions can only be bound by SUV3-a while their binding to SUV3-b is prevented by steric clashes. Thus, our results suggest that the two active sites of the SUV3 dimer are not functionally equivalent. Consistent with this, we observe RNA density only in one of the two SUV3 subunits (SUV3-a). During the preparation of this manuscript, a structure of the free SUV3 dimer was reported ^47^ which independently confirms these observations and demonstrates that the architecture of the SUV3 dimer is highly similar in its free and degradosome-bound form.

PNPase adopts a unique conformation in the degradosome which has not been observed previously. In particular, the PH2 loops are disordered, which may facilitate the transition of the KH domain to the observed outward-rotated position. This orients the conserved GXXG motif (GPGG in human PNPase) into the central channel, resulting in a narrowing of the pore. In addition, the S1 domains adopt an asymmetric arrangement which creates the binding site for SUV3. The observed conformation of PNPase positions key elements involved in RNA binding in similar positions as observed in the RNA-bound bacterial PNPase structures and creates a central channel through which RNA can enter the catalytic core from the top. Surprisingly, recent structures of ssRNA-bound human PNPase challenge the notion that RNA enters PNPase from the top. In these structures, the PH1 and PH2 loops adopt conformations that occlude the central pore and restrict access to the channel ^50^, and the PH1 loops were proposed to function as allosteric regulators, restricting catalysis to a single subunit at a time and thereby enforcing a sequential mode of activity ^50^. It was suggested that this closure redirects ssRNA toward the catalytic site via an alternative entry route from the bottom ^50^. By contrast, we observe RNA density within the entry channel at the top, which suggests that the RNA runs between the three GPGG motifs in the KH domains and then enters the catalytic core in between the three PH1 loops, which delineate the entrance to the RNA degradation chamber. Therefore, both the pore architecture and RNA density suggest that in the context of the complete degradosome complex, RNA enters PNPase from the top in a similar fashion as observed for bacterial PNPase ^58,59^.

The structure further reveals how SUV3 and PNPase interact to form the degradosome, which rationalizes a large boy of previous biochemical data. The interaction is mediated by the S1 domains of PNPase, explaining why these are required for the interaction ^46^ and why pathogenic mutations within these domains impair interactions with SUV3 while preserving RNA binding ^49^. The interaction is facilitated by the unique arrangement of the S1 domains, which create a cradle-like structure to accommodate the SUV3 dimer. This arrangement also creates a continuous positively charged path from the putative RNA exit point at SUV3-b all the way to the entry of the PNPase catalytic chamber, suggesting how RNA may be guided from the helicase to the nuclease.

Structural comparisons show that the architecture and mode of helicase-nuclease coupling markedly differ from those in yeast mtExo and the nucleo-cytoplasmic exosome. Thus, RNA degradation in human mitochondria is carried out by an architecturally unique and dedicated molecular machinery. This reflects a broader evolutionary trend towards divergence of mitochondrial gene expression machineries: just as mitochondrial transcription^62^ and RNA processing ^63,64^ have evolved unique molecular solutions distinct from their bacterial ancestors and nuclear counterparts, RNA degradation in human mitochondria is carried out by an architecturally distinct machinery. Thus, mitochondria serve as a paradigm for understanding how essential RNA metabolism pathways can be reinvented during evolution.

Finally, the structure also suggests how the substrate may be channelled from the helicase to the nuclease during processive RNA degradation. From the observed RNA densities, two different models can be envisioned: in the first, the RNA runs from SUV3-a through SUV3-b and is handed over directly to PNPase; in the second, the RNA bypasses SUV3-b and re-enters the complex at the interface between helicase and nuclease (Figure 7). As the reconstruction lacks continuous density for the RNA between SUV3-a and the helicase-nuclease interface, we cannot unambiguously distinguish between these two models at present. One possible explanation is that our cryo-EM conditions lacked ATP, which the SUV3 helicase may require to efficiently translocate along the substrate. Accordingly, we cannot formally exclude that the discontinuous RNA densities reflect more than one RNA molecule or partially occupied substrate states rather than a single continuously threaded RNA. So far, however, we have not been able to determine a structure of the complex in the presence of ATP. Further structural and biochemical studies will therefore be required to shed light on the precise path of the RNA during degradation.

Taken together, our study reveals the architecture of the human mitochondrial degradosome and provides insights into its interactions with RNA, thereby establishing a molecular and structural framework for future studies into human mitochondrial RNA degradation, its regulation, and association with disease.

## METHODS

### Molecular Cloning

Human SUV3 (residues 47–786, lacking the predicted N-terminal mitochondrial localization sequence) was amplified from human cDNA using a phosphorylated forward primer (pETSUMO_Δ46SUV3_Forward, S3) and a reverse primer that introduced a stop codon followed by a NotI restriction site (pETSUMO_ Δ46SUV3_NotI_STOP_Reverse, Extended Data Table 3). The PCR product, digested with NotI, was inserted into a modified pET-SUMO vector (a kind gift from Ricarda Richter-Dennerlein) digested with EheI and NotI. This strategy allowed cloning of SUV3 immediately downstream of the His-SUMO tag without a linker sequence, enabling the production of a scarless protein (i.e., with no extra residues at the N-terminus) following ULP1 cleavage.

All PNPase constructs were amplified from the pET15b_d23PNPase plasmid (a kind gift from Dmitry Temiakov), which encodes human PNPase (residues 24–783) in-frame with a C-terminal His tag. A variant lacking the first 39 amino acids corresponding to the predicted mitochondrial localization sequence (d39PNPase_C-His; residues 40–783 plus C-terminal His tag) was amplified using the primers 14A_LIC-v2F_Δ39PNPase_ Forward and 14A_Δ39PNPase-His_LIC-v2R_Reverse (Extended Data Table 3), and subcloned into the pET-derived 14-A vector ^65^ (Addgene plasmid #48307; a kind gift from Scott Gradia) using ligation-independent cloning (LIC) for *E. coli* expression ^65^. A variant lacking the first 23 amino acids (d23PNPase_C-His) was amplified using primers 438A_LIC-vBacF_Met_Δ23PNPase_Forward and 438A_PNPase-His_vLIC-v1rv_Reverse (Extended Data Table 3) and cloned by LIC into the 438-A vector (Addgene plasmid #55218; a kind gift from Scott Gradia) ^65^ for expression in insect cells. All constructs were transformed into *E. coli* XL1-blue (Agilent Technologies) cells for colony screening, plasmid amplification, and DNA extraction. Sequences of final expression constructs were verified by Sanger sequencing (Microsynth Seqlab).

For expression in insect cells, the 438-A_ d23PNPase _C-His construct was transformed into *E. coli* DH10αEMBacY cells (Geneva Biotech) for bacmid recombination. Bacmids were selected by blue/white screening on LB agar plates supplemented with X-gal and IPTG. Positive clones were grown in liquid culture. Bacmid DNA was isolated using an alkaline lysis protocol followed by isopropanol precipitation and was subsequently used for transfection into insect cells.

### Protein expression

d46SUV3 and d39PNPase were expressed in *E. coli* BL21 (DE3) RIL (Agilent Technologies) cells grown in LB medium supplemented with 34 ug/mL Chloramphenicol and plasmid-specific antibiotics (100 µg/mL for pETSUMO_Δ46SUV3 and 100 µg/mL Ampicilin for 14A_Δ39PNPase-His). Cultures were incubated at 37 °C, 120 rpm for 5 hours, then shifted to 18 °C, 200 rpm for 1 hour. Protein expression was induced with 200 μM IPTG, followed by incubation at 18 °C, 200 rpm for 16 hours. Cells were harvested by centrifugation at 5000 rcf, 4 °C for 20 minutes, flash-frozen in liquid nitrogen, and stored at −80 °C. d23PNPase_C-His was expressed using an insect cell expression system as previously described ^66^. In brief, recombinant bacmids were transfected into Sf9 cells (Oxford ExpressionTechnologies, 600100) cultured in ESF921 medium (Expression Technologies) using X-treme GENE 9 transfection reagent (Sigma). The resulting V0 virus was collected 4 days post-transfection and used to generate a V1 virus stock by infecting Sf21 (Expresson Systems, 94-003F) cells in suspension culture. Cultures were maintained at 27 °C with shaking for 3 days before harvesting the V1 virus, which was subsequently stored at 4 °C. For large-scale protein expression, Hi5 cells (Oxford ExpressionTechnologies, 600100) were cultured in ESF921 medium (Expression Technologies) and infected with the V1 virus. After 4 days of incubation at 27 °C with shaking, cells were harvested by centrifugation at 238 rcf for 30 min at 4 °C, washed with PBS, and stored at –80 °C until further use.

### Protein Purification

All steps were carried out at 4 °C. *E. coli* cells overexpressing d46SUV3 were resuspended in Buffer A (50 mM HEPES pH 8.0, 500 mM NaCl, 10% glycerol, 15 mM imidazole, 1 mM DTT, 1× cOmplete™ EDTA-free protease inhibitor (Roche). Cell lysis was performed by sonification and the lysate was clarified by centrifugation at 38,000 rcf, 4 °C for 1 hour. The supernatant was filtered through membranes with sieve width of 5 μm (Merck Millipore) applied to a HisTrap HP 5 mL column (Cytiva Life Sciences) pre-equilibrated with Buffer A. The column was washed with 10 column volumes (CV) of Buffer A, and bound proteins were eluted with a 0 - 100% gradient of Buffer B (50 mM HEPES pH 8.0, 500 mM NaCl, 10% glycerol, 500 mM imidazole, 1 mM DTT) over 20 CV. Elution fractions were analyzed by SDS-PAGE and fractions containing SUV3 were pooled and incubated overnight with 250 µg ULP1 protease (homemade) during dialysis in buffer C (50 mM HEPES pH 8.0, 500 mM NaCl, 10% glycerol, 1 mM DTT) to cleave the His–SUMO tag and remove imidazole. The dialyzed sample was was filtered through a 0.8 μm membrane and applied to a HisTrap HP 5 mL column equilibrated in Buffer A, followed by a 5 CV wash. The flow-through and wash fractions containing the cleaved, tag-free protein were pooled and diluted in buffer D (50 mM HEPES pH 8.0, 10% glycerol, 1 mM DTT) to reach a final NaCl concentration of 150 mM. The diluted sample was again filtered through 0.8 μm membrane and then applied to a HiTrap Heparin HP 5 mL column (Cytiva Life Sciences) pre-equilibrated with 50 mM HEPES pH 8.0, 150 mM NaCl, 10% glycerol, 1 mM DTT. Protein was eluted using a 15–100% gradient of buffer E (50 mM HEPES pH 8.0, 1 M NaCl, 10% glycerol, 1 mM DTT). Peak fractions were analyzed by SDS-PAGE. Fractions containing SUV3 were pooled, concentrated using a MWCO 30,000 Amicon Ultra Centrifugal Filter (Merck Millipore) and applied to a Superdex 200 Increase 10/300 GL column (Cytiva Life Sciences) pre-equilibrated with buffer F (20 mM HEPES pH 8.0, 300 mM NaCl, 5% glycerol, 1 mM DTT). Final peak fractions were analyzed by SDS-PAGE, pooled, concentrated using a MWCO 30,000 Amicon Ultra Centrifugal Filter (Merck), flash-frozen in liquid nitrogen, and stored at −80 °C. d39PNPase_C-His expressed in *E. coli* were purified following the same protocol, excluding the dialysis and ULP1 cleavage steps.

For purification of d23PNPase_C-His, insect cell pellets were resuspended in Buffer A, lysed by sonication, and clarified by ultracentrifugation at 235,000 rcf for 60 min at 4 °C. The soluble fraction was collected, and subsequent purification steps were performed as described for d39PNPase_C-His.

### Analytical size exclusion chromatography

Complex formation was probed by analytical size-exclusion chromatography (SEC) on an ÄKTAmicro system (GE Healthcare). Recombinant SUV3 and PNPase were mixed at a molar ratio of 2:3, with final concentrations of 20 µM and 30 µM, respectively. The mixture was incubated at room temperature for 30 min and centrifuged at 18000 rcf for 10 min at 4 °C prior to injection. Samples were loaded onto a Superdex 200 Increase 3.2/300 column (Cytiva) equilibrated in Complex assembly buffer (20 mM HEPES pH 8.0, 150 mM NaCl, 1 mM DTT) at 4 °C. Control runs with individual proteins at the same concentrations were performed under identical conditions. Chromatograms were processed and plotted using GraphPad Prism. Fractions corresponding to distinct elution peaks were collected and analyzed by SDS–PAGE.

### RNA Degradation Assays

RNA degradation assays were performed in buffer containing 20 mM HEPES (pH 8.0), 50 mM NaCl, 1 mM DTT, and 0.1 mg/mL BSA, using 100 nM of 5′-FAM-labeled random single-stranded RNA (ssRNA) or double-stranded RNA (dsRNA) oligonucleotides (IDT) as substrates (Extended Data Table 1). Sodium phosphate (NaH_2_PO_4_) and MgCl_2_ were added to final concentrations of 5 mM and 1 mM, respectively, unless otherwise stated. dsRNA substrates were prepared in water by annealing complementary RNA oligos (Extended Data Table 1), by first heating them to 95 °C, followed by stepwise cooling to 4 °C at a rate of 1 °C/min. Reactions were initiated by adding purified Δ46SUV3 and Δ39PNPT1_C-His to the RNA diluted in reaction buffer in the presence or absence of 100 μM ATP. Samples were incubated at 37 °C and reactions were stopped at various time points by the addition of 2x TBE–Urea Sample Buffer (Sigma). Control reactions consisted of RNA-only samples incubated under identical conditions. Proteins in the samples were digested with Proteinase K (NEB) at 50 °C for 30 minutes, followed by heat denaturation at 95 °C for 5 minutes. Samples were separated on a 20% TBE–urea polyacrylamide gel at 300 V for 90 minutes. Degradation products were visualized using a Typhoon FLA 9500 scanner (GE Healthcare). RNA fragment lengths were estimated by comparison to a ladder of synthesized FAM-labeled ssRNA molecules of known sizes.

### Cryo-EM sample preparation and data collection

The degradosome complex was assembled by mixing 3 µM PNPase (d23PNPase_C–His) with 6 µM SUV3 (d46SUV3) in a final buffer containing 25 mM Tris-HCl (pH 8.0), 70 mM NaCl, 1% (v/v) glycerol, 1 mM dithiothreitol (DTT), and 5 mM sodium phosphate (NaH_2_PO_4_). The substrate RNA (dsRNA_02_PT, 6 µM) was added and the sample was incubated at 37 °C for 1 hour, then kept on ice until grid preparation. 4 µL of sample were applied to freshly glow-discharged Quantifoil R 2/1 holey carbon grids (Quantifoil) at 4 °C and 95% humidity. Grids were blotted for 5 seconds with a blot force of 5 using a Vitrobot Mark IV (Thermo Fisher Scientific) and subsequently plunge-frozen in liquid ethane.

Cryo-EM data were collected using a Titan Krios transmission electron microscope (Thermo Fisher Scientific) operated at 300 keV and controlled with SerialEM ^67,68^. Images were acquired in energy-filtered transmission electron microscopy (EFTEM) mode using a GIF quantum energy filter with a slit width of 20 eV, and a K3 direct electron detector (Gatan) at a nominal magnification of 105,000×, corresponding to a calibrated pixel size of 0.834 Å/pixel. Exposures were recorded in counting mode for 2.186 s at a dose rate of 15.91e^−^/pixel/s, yielding a total dose of 50 e^−^/Å^2^, fractionated into 50 movie frames. Images were acquired in a 3×3-hole pattern per stage movement, with active beam tilt compensation implemented via SerialEM.

### Cryo-EM data processing and analysis

Motion correction, CTF-estimation and particle picking were performed on the fly using Warp version 1.0.9 ^69^. Further processing was carried out using cryoSPARC v.4.5.3 ^70^ and RELION-5 ^71^. A representative micrograph and the cryo-EM processing workflow are depicted in Extended Data Figure 2a-b. and S2B. For initial processing steps, the datasets were split into batches and particles were extracted in RELION with a box size of 448 px with 2x binning. Initial 2D classification was done in Cryosparc. To obtain an initial reconstruction, particles sorted into degradosome classes were selected and used for ab initio reconstruction followed by non-uniform refinement in Cryosparc. The initial volume was then used for supervised classification of the entire dataset in 3D. For this, the initial degradosome volume was used as an input model for heterogeneous refinement in CryoSPARC, alongside five “junk” models generated *ab initio* from poor particles selected in 2D classification. Classification of the entire dataset using this strategy yielded a subset of 3.2 million particles assigned to the degradosome class. These particles were subjected to unsupervised classification, reducing the set to 615,000 particles with improved SUV3 occupancy. This subset was then subjected to one round of unsupervised 3D classification in RELION. Particles from the best-resolved classes (468,000 particles) were re-extracted without binning, 3D refined and subjected to a final round of unsupervised classification. The best-aligning classes (210,000 particles) were subsequently refined in 3D. Post-processing was performed to generate half-maps for CTF refinement. Following CTF refinement, an additional round of 3D refinement yielded a map at 3.8 Å resolution, which improved to 3.3 Å after post-processing (Map A, consensus map).

To improve SUV3 resolution, multibody refinement ^53^ with blush regularization ^72^ was carried out using two masks, one encompassing PNPase, excluding the S1 domains (body 1), and another including SUV3 and S1 domains of PNPase (body 2).Following post-processing of the individual bodies obtained from multibody refinement, local resolutions of 3.2 Å for the PNPase region (Map B) and 3.6 Å for the SUV3 region (Map C) were achieved. The focused maps were combined into a composite map (Map D) using the vopmaximum command in UCSF ChimeraX ^73,74^.

### Model building and refinement

Initial model building was performed by rigid-body fitting two copies of the SUV3 crystal structure (PDB: 3RC8 ^23^) and the cryo-EM structure of human PNPase in its open conformation (PDB: 9KJR ^49^) into the cryo-EM maps using UCSF ChimeraX ^73,74^. The S1 domains were omitted during the initial fitting and were instead docked separately into the density as individual rigid bodies. The resulting model was manually rebuilt and refined in Coot 0.9.8.96 ^75^. Following this, an additional density was observed in the active site of SUV3-a monomer. A 4-nt poly(U) RNA fragment was modeled into this density, guided by the SUV3–ssRNA co-crystal structure (PDB: 3RC8 ^23^). Additional regions of unmodeled, discontinuous and less well-resolved density were observed, which we hypothesize to correspond to RNA. However, due to the insufficient resolution in these areas, we refrained from modelling any further RNA. Initial refinement was performed using molecular dynamics–assisted refinement in ISOLDE ^76^ with hydrogens included, followed by real-space refinement in Phenix ^77,78^, using the masked, post-processed composite EM map (Map A). The final model was validated using the MolProbity package within the Phenix suite ^79^. Refinement statistics are provided in Table S2. All structural analyses and figure generation were carried out using UCSF ChimeraX ^73,74^.

## DATA AVAILABILITY

The UniProt accession numbers for PNPase and SUV3 are Q8TCS8 and Q8IYB8. The electron potential reconstructions were deposited with the Electron Microscopy Database (EMDB) under accession codes: EMD-58691 (Map A: Consensus Map - Structure of the human mitochondrial RNA degradosome), EMD-58689 (Map B: Focused refinement - PNPase lacking S1 domains), EMD-58690 (Map C: Focused refinement - SUV3 dimer and S1 domains of PNPase) and EMD-58688 (Map D - Composite Map - Structure of the human mitochondrial RNA degradosome complex).The structure coordinates were deposited to the Protein Data Bank (PDB) under accession code 31WW (Structure of the human mitochondrial RNA degradosome complex). Source data are provided with this paper.

## ACKNOWLEDGEMENTS

We thank all members of the Hillen Lab for helpful discussions, in particular Katja Ditter for support and training in insect cell culture. We thank Christian Dienemann and Ulrich Steuerwald (MPI-NAT cryo-EM facility) for assistance with cryo-EM data acquisition.

H.S.H was supported by the Deutsche Forschungsgemeinschaft under Germany’s Excellence Strategy EXC 2067/1-390729940 and SFB1565 (Project number 469281184, P13), by the European Union (ERC Starting Grant MitoRNA, grant agreement no. 101116869) and by the EMBO Young Investigator Programme. P.F.V.d.P was supported by Germany’s Excellence Strategy EXC 2067/1-390729940 and Hertha Sponer College. Views and opinions expressed are however those of the author(s) only and do not necessarily reflect those of the European Union or the European Research Council Executive Agency. Neither the European Union nor the granting authority can be held responsible for them.

## AUTHOR CONTRIBUTIONS

H.S.H. designed and supervised the research. P.F.V.d.P. cloned, expressed, and established protein purification protocols and biochemical assays, prepared cryo-EM grids, collected cryo-EM data and carried our image processing and model building. A.B. assisted in cryo-EM data processing. S.S., A.V., M.P.M., A.P.L.D. assisted with protein purification and biochemical assays. P.F.V.d.P., B.K. and H.S.H interpreted the data and wrote the manuscript, with input from all authors.

## STATEMENT ON USE OF AI

During the preparation of this work, the authors used ChatGPT-5.5 in order to check for grammar and improve the readability of parts of the manuscript. After using this tool/service, the authors reviewed and edited the content as needed and take full responsibility for the content of the published article.

Correspondence should be addressed to

## REFERENCES

1. Anderson, S., Bankier, A.T., Barrell, B.G., Bruijn, M.H.L. de, Coulson, A.R., Drouin, J., Eperon, I.C., Nierlich, D.P., Roe, B.A., Sanger, F., et al. (1981). Sequence and organization of the human mitochondrial genome. Nature 290, 457–465. 10.1038/290457a0.

2. Ojala, D., Montoya, J., and Attardi, G. (1981). tRNA punctuation model of RNA processing in human mitochondria. Nature 290, 470–474. 10.1038/290470a0.

3. Crews, S., Ojala, D., Posakony, J., Nishiguchi, J., and Attardi, G. (1979). Nucleotide sequence of a region of human mitochondrial DNA containing the precisely identified origin of replication. Nature 277, 192–198. 10.1038/277192a0.

4. Barchiesi, A., and Vascotto, C. (2019). Transcription, Processing, and Decay of Mitochondrial RNA in Health and Disease. Int. J. Mol. Sci. 20, 2221. 10.3390/ijms20092221.

5. Hällberg, B.M., and Larsson, N.-G. (2014). Making Proteins in the Powerhouse. Cell Metab. 20, 226–240. 10.1016/j.cmet.2014.07.001.

6. Falkenberg, M., Larsson, N.-G., and Gustafsson, C.M. (2007). DNA Replication and Transcription in Mammalian Mitochondria. Annu. Rev. Biochem. 76, 679–699. 10.1146/annurev.biochem.76.060305.152028.

7. Chujo, T., Ohira, T., Sakaguchi, Y., Goshima, N., Nomura, N., Nagao, A., and Suzuki, T. (2012). LRPPRC/SLIRP suppresses PNPase-mediated mRNA decay and promotes polyadenylation in human mitochondria. Nucleic Acids Res. 40, 8033–8047. 10.1093/nar/gks506.

8. Rackham, O., Mercer, T.R., and Filipovska, A. (2012). The human mitochondrial transcriptome and the RNA-binding proteins that regulate its expression. Wiley Interdiscip. Rev.: RNA 3, 675–695. 10.1002/wrna.1128.

9. Pearce, S.F., Rebelo-Guiomar, P., D’Souza, A.R., Powell, C.A., Haute, L.V., and Minczuk, M. (2017). Regulation of Mammalian Mitochondrial Gene Expression: Recent Advances. Trends Biochem. Sci. 42, 625–639. 10.1016/j.tibs.2017.02.003.

10. Zhu, X., Xie, X., Das, H., Tan, B.G., Shi, Y., Al-Behadili, A., Peter, B., Motori, E., Valenzuela, S., Posse, V., et al. (2022). Non-coding 7S RNA inhibits transcription via mitochondrial RNA polymerase dimerization. Cell 185, 2309–2323.e24. 10.1016/j.cell.2022.05.006.

11. Silva, S., Camino, L.P., and Aguilera, A. (2018). Human mitochondrial degradosome prevents harmful mitochondrial R loops and mitochondrial genome instability. Proc. Natl. Acad. Sci. 115, 11024–11029. 10.1073/pnas.1807258115.

12. Grochowska, J., Czerwinska, J., Borowski, L.S., and Szczesny, R.J. (2022). Mitochondrial RNA, a new trigger of the innate immune system. Wiley Interdiscip. Rev.: RNA 13, e1690. 10.1002/wrna.1690.

13. Lin-Chao, S., Chiou, N.-T., and Schuster, G. (2007). The PNPase, exosome and RNA helicases as the building components of evolutionarily-conserved RNA degradation machines. J. Biomed. Sci. 14, 523–532. 10.1007/s11373-007-9178-y.

14. Coburn, G.A., Miao, X., Briant, D.J., and Mackie, G.A. (1999). Reconstitution of a minimal RNA degradosome demonstrates functional coordination between a 3′ exonuclease and a DEAD-box RNA helicase. Genes Dev. 13, 2594–2603. 10.1101/gad.13.19.2594.

15. Januszyk, K., and Lima, C.D. (2014). The eukaryotic RNA exosome. Curr. Opin. Struct. Biol. 24, 132–140. 10.1016/j.sbi.2014.01.011.

16. Büttner, K., Wenig, K., and Hopfner, K. (2006). The exosome: a macromolecular cage for controlled RNA degradation. Mol. Microbiol. 61, 1372–1379. 10.1111/j.1365-2958.2006.05331.x.

17. Carpousis, A.J., Houwe, G.V., Ehretsmann, C., and Krisch, H.M. (1994). Copurification of E. coli RNAase E and PNPase: Evidence for a specific association between two enzymes important in RNA processing and degradation. Cell 76, 889–900. 10.1016/0092-8674(94)90363-8.

18. Carpousis, A.J. (2002). The Escherichia coli RNA degradosome: structure, function and relationship to other ribonucleolytic multienyzme complexes. Biochem. Soc. Trans. 30, 150–155. 10.1042/bst0300150.

19. Carpousis, A.J. (2007). The RNA Degradosome of Escherichia coli: An mRNA-Degrading Machine Assembled on RNase E. Annu. Rev. Microbiol. 61, 71–87. 10.1146/annurev.micro.61.080706.093440.

20. Py, B., Causton, H., Mudd, E.A., and Higgins, C.F. (1994). A protein complex mediating mRNA degradation in Escherichia coli. Mol. Microbiol. 14, 717–729. 10.1111/j.1365-2958.1994.tb01309.x.

21. Hardwick, S.W., and Luisi, B.F. (2013). Rarely at rest. RNA Biol. 10, 56–70. 10.4161/rna.22270.

22. Razew, M., Warkocki, Z., Taube, M., Kolondra, A., Czarnocki-Cieciura, M., Nowak, E., Labedzka-Dmoch, K., Kawinska, A., Piatkowski, J., Golik, P., et al. (2018). Structural analysis of mtEXO mitochondrial RNA degradosome reveals tight coupling of nuclease and helicase components. Nat. Commun. 9, 97. 10.1038/s41467-017-02570-5.

23. Jedrzejczak, R., Wang, J., Dauter, M., Szczesny, R.J., Stepien, P.P., and Dauter, Z. (2011). Human Suv3 protein reveals unique features among SF2 helicases. Acta Crystallogr. Sect. D 67, 988–996. 10.1107/s0907444911040248.

24. Shu, Z., Vijayakumar, S., Chen, C.-F., Chen, P.-L., and Lee, W.-H. (2004). Purified Human SUV3p Exhibits Multiple-Substrate Unwinding Activity upon Conformational Change †. Biochemistry 43, 4781–4790. 10.1021/bi0356449.

25. Szczesny, R.J., Wojcik, M.A., Borowski, L.S., Szewczyk, M.J., Skrok, M.M., Golik, P., and Stepien, P.P. (2013). Yeast and human mitochondrial helicases. Biochim. Biophys. Acta (BBA) - Gene Regul. Mech. 1829, 842–853. 10.1016/j.bbagrm.2013.02.009.

26. Falchi, F.A., Pizzoccheri, R., and Briani, F. (2022). Activity and Function in Human Cells of the Evolutionary Conserved Exonuclease Polynucleotide Phosphorylase. Int. J. Mol. Sci. 23, 1652. 10.3390/ijms23031652.

27. Wang, D.D.-H., Shu, Z., Lieser, S.A., Chen, P.-L., and Lee, W.-H. (2009). Human Mitochondrial SUV3 and Polynucleotide Phosphorylase Form a 330-kDa Heteropentamer to Cooperatively Degrade Double-stranded RNA with a 3′-to-5′ Directionality*. J. Biol. Chem. 284, 20812–20821. 10.1074/jbc.m109.009605.

28. Pajak, A., Laine, I., Clemente, P., El-Fissi, N., Schober, F.A., Maffezzini, C., Calvo-Garrido, J., Wibom, R., Filograna, R., Dhir, A., et al. (2019). Defects of mitochondrial RNA turnover lead to the accumulation of double-stranded RNA in vivo. PLoS Genet. 15, e1008240. 10.1371/journal.pgen.1008240.

29. Dhir, A., Dhir, S., Borowski, L.S., Jimenez, L., Teitell, M., Rötig, A., Crow, Y.J., Rice, G.I., Duffy, D., Tamby, C., et al. (2018). Mitochondrial double-stranded RNA triggers antiviral signalling in humans. Nature 560, 238–242. 10.1038/s41586-018-0363-0.

30. von Ameln, S., Wang, G., Boulouiz, R., Rutherford, M.A., Smith, G.M., Li, Y., Pogoda, H.-M., Nürnberg, G., Stiller, B., Volk, A.E., et al. (2012). A Mutation in PNPT1, Encoding Mitochondrial-RNA-Import Protein PNPase, Causes Hereditary Hearing Loss. Am. J. Hum. Genet. 91, 919–927. 10.1016/j.ajhg.2012.09.002.

31. Li, Y.-Y., Gao, Y., Zhong, X.-X., and Chen, G.-F. (2025). A novel polyribonucleotide nucleotidyltransferase 1 (PNPT1) gene variant potentially associated with combined oxidative phosphorylation deficiency 13: case report and literature review. Transl. Pediatr. 14, 33849–33349. 10.21037/tp-24-419.

32. Rius, R., Bergen, N.J.V., Compton, A.G., Riley, L.G., Kava, M.P., Balasubramaniam, S., Amor, D.J., Fanjul-Fernandez, M., Cowley, M.J., Fahey, M.C., et al. (2019). Clinical Spectrum and Functional Consequences Associated with Bi-Allelic Pathogenic PNPT1 Variants. J. Clin. Med. 8, 2020. 10.3390/jcm8112020.

33. Matilainen, S., Carroll, C.J., Richter, U., Euro, L., Pohjanpelto, M., Paetau, A., Isohanni, P., and Suomalainen, A. (2017). Defective mitochondrial RNA processing due to PNPT1 variants causes Leigh syndrome. Hum. Mol. Genet. 26, 3352–3361. 10.1093/hmg/ddx221.

34. Haddad, S., Record, C.J., Self, E., Skorupinska, M., Rossor, A.M., Laura, M., Ingle, G., Manzur, A., Muntoni, F., Blake, J.C., et al. (2025). Heterozygous PNPT1 Variants Cause a Sensory Ataxic Neuropathy. Eur. J. Neurol. 32, e70064. 10.1111/ene.70064.

35. Barbier, M., Bahlo, M., Pennisi, A., Jacoupy, M., Tankard, R.M., Ewenczyk, C., Davies, K.C., Lino-Coulon, P., Colace, C., Rafehi, H., et al. (2022). Heterozygous PNPT1 Variants Cause Spinocerebellar Ataxia Type 25. Ann. Neurol. 92, 122–137. 10.1002/ana.26366.

36. Eaton, A., Bernier, F.P., Goedhart, C., Caluseriu, O., Lamont, R.E., Boycott, K.M., Parboosingh, J.S., Innes, A.M., and Consortium, C.C. (2018). Is PNPT1-related hearing loss ever non-syndromic? Whole exome sequencing of adult siblings expands the natural history of PNPT1-related disorders. Am. J. Méd. Genet. Part A 176, 2487–2493. 10.1002/ajmg.a.40516.

37. Vedrenne, V., Gowher, A., De Lonlay, P., Nitschke, P., Serre, V., Boddaert, N., Altuzarra, C., Mager-Heckel, A.-M., Chretien, F., Entelis, N., et al. (2012). Mutation in PNPT1, which Encodes a Polyribonucleotide Nucleotidyltransferase, Impairs RNA Import into Mitochondria and Causes Respiratory-Chain Deficiency. Am. J. Hum. Genet. 91, 912–918. 10.1016/j.ajhg.2012.09.001.

38. Sato, R., Arai-Ichinoi, N., Kikuchi, A., Matsuhashi, T., Numata-Uematsu, Y., Uematsu, M., Fujii, Y., Murayama, K., Ohtake, A., Abe, T., et al. (2018). Novel biallelic mutations in the PNPT1 gene encoding a mitochondrial-RNA-import protein PNPase cause delayed myelination. Clin. Genet. 93, 242–247. 10.1111/cge.13068.

39. Kuht, H.J., Thomas, K.A., Hisaund, M., Maconachie, G.D.E., and Thomas, M.G. (2021). Ocular Manifestations of PNPT1-Related Neuropathy. J. Neuro-Ophthalmol. 41, e293–e296. 10.1097/wno.0000000000001012.

40. Bamborschke, D., Kreutzer, M., Koy, A., Koerber, F., Lucas, N., Huenseler, C., Herkenrath, P., Lee-Kirsch, M.A., and Cirak, S. (2021). PNPT1 mutations may cause Aicardi-Goutières-Syndrome. Brain Dev. 43, 320–324. 10.1016/j.braindev.2020.10.005.

41. Sgobbi, P., Farias, I.B., Serrano, P. de L., Badia, B. de M.L., Oliveira, H.B. de, Barbosa, A.S., Pereira, C.A., Moreira, V. de F., Chieia, M.A.T., Barbosa, A.R., et al. (2024). PNPT1 Spectrum Disorders: An Underrecognized and Complex Group of Neurometabolic Disorders. Muscles 3, 4–15. 10.3390/muscles3010002.

42. Tenorio, R.B., Lira, J.S.S. de, Cordellini, M.F., and Donis, K.C. (2025). Unveiling Spinocerebellar Ataxia 25: First Case Report of a Brazilian Family. Cerebellum 24, 41. 10.1007/s12311-025-01794-2.

43. Alodaib, A., Sobreira, N., Gold, W.A., Riley, L.G., Bergen, N.J.V., Wilson, M.J., Bennetts, B., Thorburn, D.R., Boehm, C., and Christodoulou, J. (2017). Whole-exome sequencing identifies novel variants in PNPT1 causing oxidative phosphorylation defects and severe multisystem disease. Eur. J. Hum. Genet. 25, 79–84. 10.1038/ejhg.2016.128.

44. Esveld, S.L. van, Rodenburg, R.J., Al-Murshedi, F., Al-Ajmi, E., Al-Zuhaibi, S., Huynen, M.A., and Spelbrink, J.N. (2022). Mitochondrial RNA processing defect caused by a SUPV3L1 mutation in two siblings with a novel neurodegenerative syndrome. J. Inherit. Metab. Dis. 45, 292–307. 10.1002/jimd.12476.

45. Tsygankova, P., Chistol, D., Krylova, T., Bychkov, I., Tabakov, V., Markova, T., Dadali, E., and Zakharova, E. (2024). A New Case of Mitochondrial RNA Helicase SUPV3L1-Associated Neurodegenerative Disease: Ataxia, Spasticity, Optic Atrophy, and Skin Hypopigmentation (ASOASH). Genes 15, 1406. 10.3390/genes15111406.

46. Jain, M., Golzarroshan, B., Lin, C., Agrawal, S., Tang, W., Wu, C., and Yuan, H.S. (2022). Dimeric assembly of human Suv3 helicase promotes its RNA unwinding function in mitochondrial RNA degradosome for RNA decay. Protein Sci. 31, e4312. 10.1002/pro.4312.

47. Patra, M., Jain, M., Li, Y.-C., Chen, Y.-P., Golzarroshan, B., and Yuan, H.S. (2026). Asymmetric dimeric assembly of Suv3 helicase facilitates processive RNA unwinding. Nat. Commun. 10.1038/s41467-026-71901-2.

48. Lin, C.L., Wang, Y.-T., Yang, W.-Z., Hsiao, Y.-Y., and Yuan, H.S. (2012). Crystal structure of human polynucleotide phosphorylase: insights into its domain function in RNA binding and degradation. Nucleic Acids Res. 40, 4146–4157. 10.1093/nar/gkr1281.

49. Li, Y.-C., Wang, C.-H., Patra, M., Chen, Y.-P., Yang, W.-Z., and Yuan, H.S. (2025). Structural insights into human PNPase in health and disease. Nucleic Acids Res. 53, gkaf119. 10.1093/nar/gkaf119.

50. Unseld, O., Das, H., and Hällberg, B.M. (2025). Loop-mediated regulation and base flipping drive RNA cleavage by human mitochondrial PNPase. Nucleic Acids Res. 53, gkaf1296. 10.1093/nar/gkaf1296.

51. Warnecke, J.M., Fürste, J.P., Hardt, W.D., Erdmann, V.A., and Hartmann, R.K. (1996). Ribonuclease P (RNase P) RNA is converted to a Cd(2+)-ribozyme by a single Rp-phosphorothioate modification in the precursor tRNA at the RNase P cleavage site. Proc. Natl. Acad. Sci. 93, 8924–8928. 10.1073/pnas.93.17.8924.

52. Teramoto, T., Koyasu, T., Yokogawa, T., Adachi, N., Mayanagi, K., Nakamura, T., Senda, T., and Kakuta, Y. (2025). Structural basis of transfer RNA processing by bacterial minimal RNase P. Nat. Commun. 16, 5456. 10.1038/s41467-025-60002-1.

53. Nakane, T., and Scheres, S.H.W. (2020). cryoEM, Methods and Protocols. Methods Mol. Biol. 2215, 145–160. 10.1007/978-1-0716-0966-8_7.

54. Jedrzejczak, R., Wang, J., Dauter, M., Szczesny, R.J., Stepien, P.P., and Dauter, Z. (2011). Human Suv3 protein reveals unique features among SF2 helicases. Acta Crystallogr Sect D Biological Crystallogr 67, 988–996. 10.1107/s0907444911040248.

55. Abramson, J., Adler, J., Dunger, J., Evans, R., Green, T., Pritzel, A., Ronneberger, O., Willmore, L., Ballard, A.J., Bambrick, J., et al. (2024). Accurate structure prediction of biomolecular interactions with AlphaFold 3. Nature 630, 493–500. 10.1038/s41586-024-07487-w.

56. Kempen, M. van, Kim, S.S., Tumescheit, C., Mirdita, M., Lee, J., Gilchrist, C.L.M., Söding, J., and Steinegger, M. (2024). Fast and accurate protein structure search with Foldseek. Nat. Biotechnol. 42, 243–246. 10.1038/s41587-023-01773-0.

57. Büttner, K., Nehring, S., and Hopfner, K.-P. (2007). Structural basis for DNA duplex separation by a superfamily-2 helicase. Nat. Struct. Mol. Biol. 14, 647–652. 10.1038/nsmb1246.

58. Dendooven, T., Sinha, D., Roeselová, A., Cameron, T.A., Lay, N.R.D., Luisi, B.F., and Bandyra, K.J. (2021). A cooperative PNPase-Hfq-RNA carrier complex facilitates bacterial riboregulation. Mol. Cell 81, 2901–2913.e5. 10.1016/j.molcel.2021.05.032.

59. Hardwick, S.W., Gubbey, T., Hug, I., Jenal, U., and Luisi, B.F. (2012). Crystal structure of Caulobacter crescentus polynucleotide phosphorylase reveals a mechanism of RNA substrate channelling and RNA degradosome assembly. Open Biol. 2, 120028. 10.1098/rsob.120028.

60. Nurmohamed, S., Vaidialingam, B., Callaghan, A.J., and Luisi, B.F. (2009). Crystal Structure of Escherichia coli Polynucleotide Phosphorylase Core Bound to RNase E, RNA and Manganese: Implications for Catalytic Mechanism and RNA Degradosome Assembly. J. Mol. Biol. 389, 17–33. 10.1016/j.jmb.2009.03.051.

61. Weick, E.-M., Puno, M.R., Januszyk, K., Zinder, J.C., DiMattia, M.A., and Lima, C.D. (2018). Helicase-Dependent RNA Decay Illuminated by a Cryo-EM Structure of a Human Nuclear RNA Exosome-MTR4 Complex. Cell 173, 1663–1677.e21. 10.1016/j.cell.2018.05.041.

62. Hillen, H.S., Temiakov, D., and Cramer, P. (2018). Structural basis of mitochondrial transcription. Nat. Struct. Mol. Biol. 25, 754–765. 10.1038/s41594-018-0122-9.

63. Bhatta, A., Kuhle, B., Yu, R.D., Spanaus, L., Ditter, K., Bohnsack, K.E., and Hillen, H.S. (2025). Molecular basis of human nuclear and mitochondrial tRNA 3′ processing. Nat. Struct. Mol. Biol. 32, 613–624. 10.1038/s41594-024-01445-w.

64. Bhatta, A., Dienemann, C., Cramer, P., and Hillen, H.S. (2021). Structural basis of RNA processing by human mitochondrial RNase P. Nat. Struct. Mol. Biol. 28, 713–723. 10.1038/s41594-021-00637-y.

65. Gradia, S.D., Ishida, J.P., Tsai, M.-S., Jeans, C., Tainer, J.A., and Fuss, J.O. (2017). Chapter One MacroBac: New Technologies for Robust and Efficient Large-Scale Production of Recombinant Multiprotein Complexes. Methods Enzym. 592, 1–26. 10.1016/bs.mie.2017.03.008.

66. Berger, I., Fitzgerald, D.J., and Richmond, T.J. (2004). Baculovirus expression system for heterologous multiprotein complexes. Nat. Biotechnol. 22, 1583–1587. 10.1038/nbt1036.

67. Mastronarde, D.N. (2018). Advanced Data Acquisition From Electron Microscopes With SerialEM. Microsc. Microanal. 24, 864–865. 10.1017/s1431927618004816.

68. Mastronarde, D.N. (2003). SerialEM: A Program for Automated Tilt Series Acquisition on Tecnai Microscopes Using Prediction of Specimen Position. Microsc. Microanal. 9, 1182–1183. 10.1017/s1431927603445911.

69. Tegunov, D., and Cramer, P. (2019). Real-time cryo-electron microscopy data preprocessing with Warp. Nat. Methods 16, 1146–1152. 10.1038/s41592-019-0580-y.

70. Punjani, A., Rubinstein, J.L., Fleet, D.J., and Brubaker, M.A. (2017). cryoSPARC: algorithms for rapid unsupervised cryo-EM structure determination. Nat. Methods 14, 290–296. 10.1038/nmeth.4169.

71. Burt, A., Toader, B., Warshamanage, R., Kügelgen, A. von, Pyle, E., Zivanov, J., Kimanius, D., Bharat, T.A.M., and Scheres, S.H.W. (2024). An image processing pipeline for electron cryo-tomography in RELION-5. FEBS Open Bio 14, 1788–1804. 10.1002/2211-5463.13873.

72. Kimanius, D., Jamali, K., Wilkinson, M.E., Lövestam, S., Velazhahan, V., Nakane, T., and Scheres, S.H.W. (2024). Data-driven regularization lowers the size barrier of cryo-EM structure determination. Nat. Methods 21, 1216–1221. 10.1038/s41592-024-02304-8.

73. Meng, E.C., Goddard, T.D., Pettersen, E.F., Couch, G.S., Pearson, Z.J., Morris, J.H., and Ferrin, T.E. (2023). UCSF ChimeraX : Tools for structure building and analysis. Protein Sci. 32, e4792. 10.1002/pro.4792.

74. Pettersen, E.F., Goddard, T.D., Huang, C.C., Meng, E.C., Couch, G.S., Croll, T.I., Morris, J.H., and Ferrin, T.E. (2021). UCSF ChimeraX: Structure visualization for researchers, educators, and developers. Protein Sci. 30, 70–82. 10.1002/pro.3943.

75. Casañal, A., Lohkamp, B., and Emsley, P. (2020). Current developments in Coot for macromolecular model building of Electron Cryo-microscopy and Crystallographic Data. Protein Sci. : A Publ. Protein Soc. 29, 1069–1078. 10.1002/pro.3791.

76. Croll, T.I. (2018). ISOLDE: a physically realistic environment for model building into low-resolution electron-density maps. Acta Crystallogr. Sect. D: Struct. Biol. 74, 519–530. 10.1107/s2059798318002425.

77. Adams, P.D., Afonine, P.V., Bunkóczi, G., Chen, V.B., Davis, I.W., Echols, N., Headd, J.J., Hung, L., Kapral, G.J., Grosse-Kunstleve, R.W., et al. (2010). PHENIX: a comprehensive Python-based system for macromolecular structure solution. Acta Crystallogr. Sect. D 66, 213–221. 10.1107/s0907444909052925.

78. Afonine, P.V., Poon, B.K., Read, R.J., Sobolev, O.V., Terwilliger, T.C., Urzhumtsev, A., and Adams, P.D. (2018). Real-space refinement in PHENIX for cryo-EM and crystallography. Acta Crystallogr. Sect. D 74, 531–544. 10.1107/s2059798318006551.

79. Williams, C.J., Headd, J.J., Moriarty, N.W., Prisant, M.G., Videau, L.L., Deis, L.N., Verma, V., Keedy, D.A., Hintze, B.J., Chen, V.B., et al. (2018). MolProbity: More and better reference data for improved all-atom structure validation. Protein Sci. 27, 293–315. 10.1002/pro.3330.

