## Supplemental Information for "Structure of the human mitochondrial RNA degradosome reveals a distinct mode of helicase–nuclease coupling"

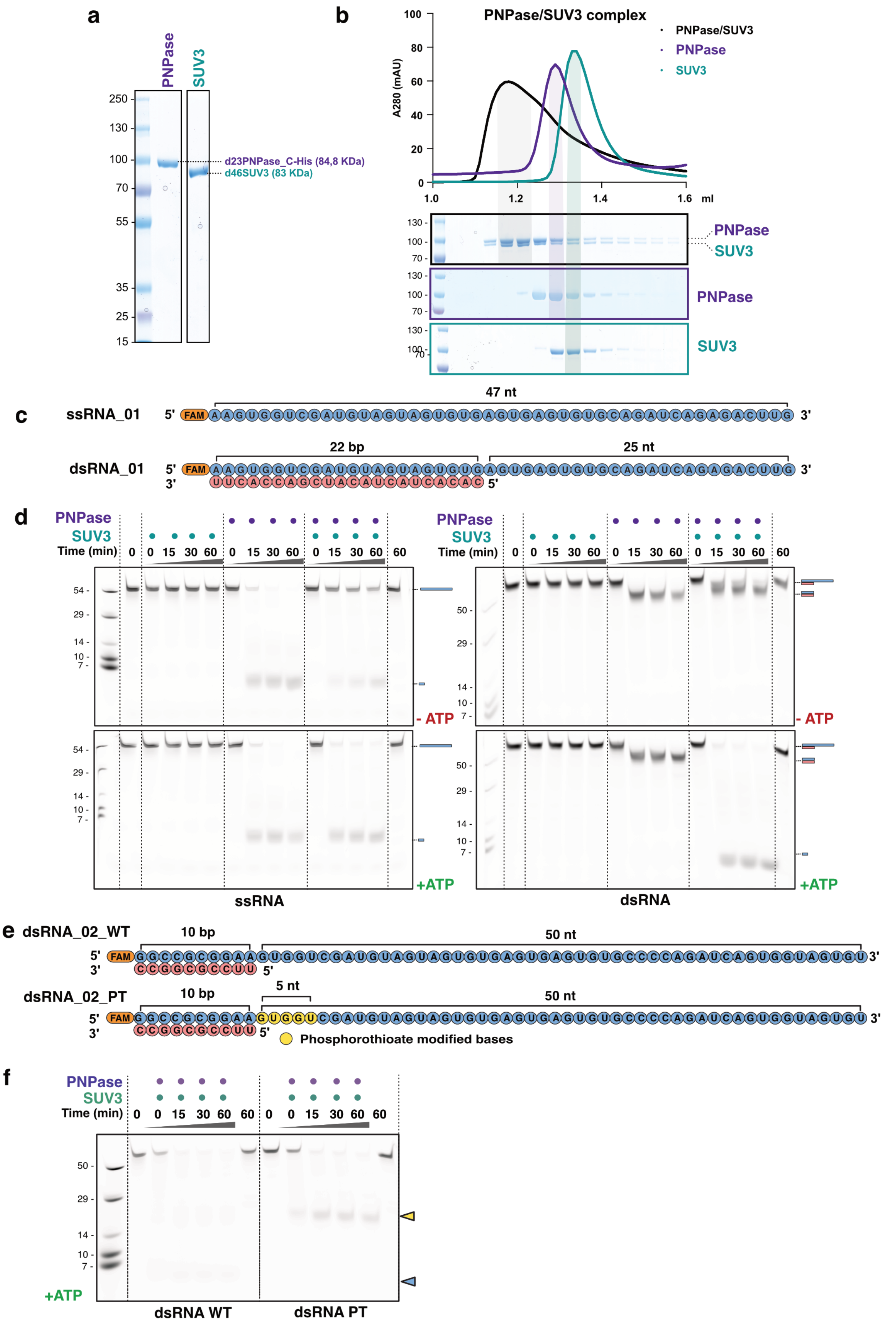


**Extended Data Figure 1 - SUV3 and PNPase assemble into a stable, catalytically active complex**

Figure captions follow on the next page.

**Extended Data Figure 1 - SUV3 and PNPase assemble into a stable, catalytically active complex**
**a)** SDS-PAGE analysis of purified PNPase and SUV3.

**b)** Analytical size-exclusion chromatography (SEC) followed by SDS-PAGE, showing that SUV3 and PNPase co-elute as a stable complex.

**c)** Schematic representation of the single-stranded RNA (ssRNA_01) and double-stranded RNA (dsRNA_01) scaffolds used to assay complex activity.

**d)** RNA degradation activity assay performed using purified proteins. Complete degradation of dsRNA is observed only in the presence of PNPase, SUV3, and ATP, indicating that all three components are required for dsRNA degradation. Gels are representative of three technical replicates.

**e)** Schematic representation of the double-stranded RNA (dsRNA_02_PT) scaffold used for cryo-EM analysis. PT indicates phosphorothioate modifications. An equivalent probe lacking phosphorothioate modifications (dsRNA_02_WT) was used in activity assays to demonstrate that the PT modifications prevent complete degradation of the substrate.

**f)** RNA degradation assay using cryo-EM substrate. In the presence of ATP, the SUV3–PNPase complex degrades the WT substrate completely (blue arrow). Degradation of the PT-modified substrate results in a larger intermediate (yellow arrow), indicating stalling of the degradation. Gels are representative of three technical replicates.

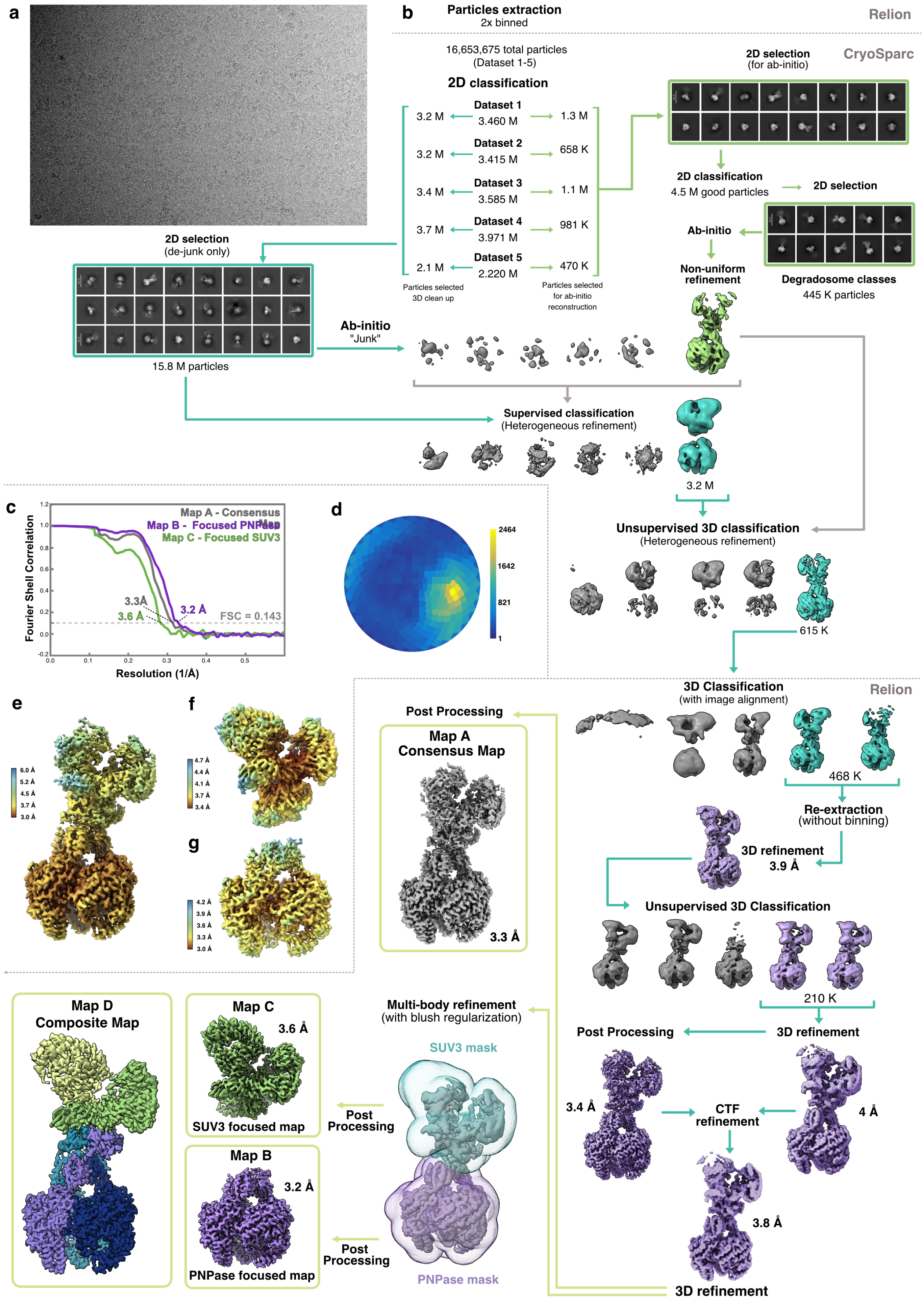


**Extended Data Figure 2 - Cryo-EM processing workflow**

Figure captions follow on the next page.

**Extended Data Figure 2 - Cryo-EM processing workflow**

**a)** Exemplary cryo-EM micrograph of the degradosome dataset.

**b)** Cryo-EM processing workflow for structure determination.

**c)** Fourier Shell Correlation (FSC)-plots for cryo-EM reconstructions used.

**d)** Angular distribution plot of Map A. Created with Warp [1].

**e)** Local resolution map of Map A.

**f)** Local resolution map of Map C.

**g)** Local resolution map of Map B.

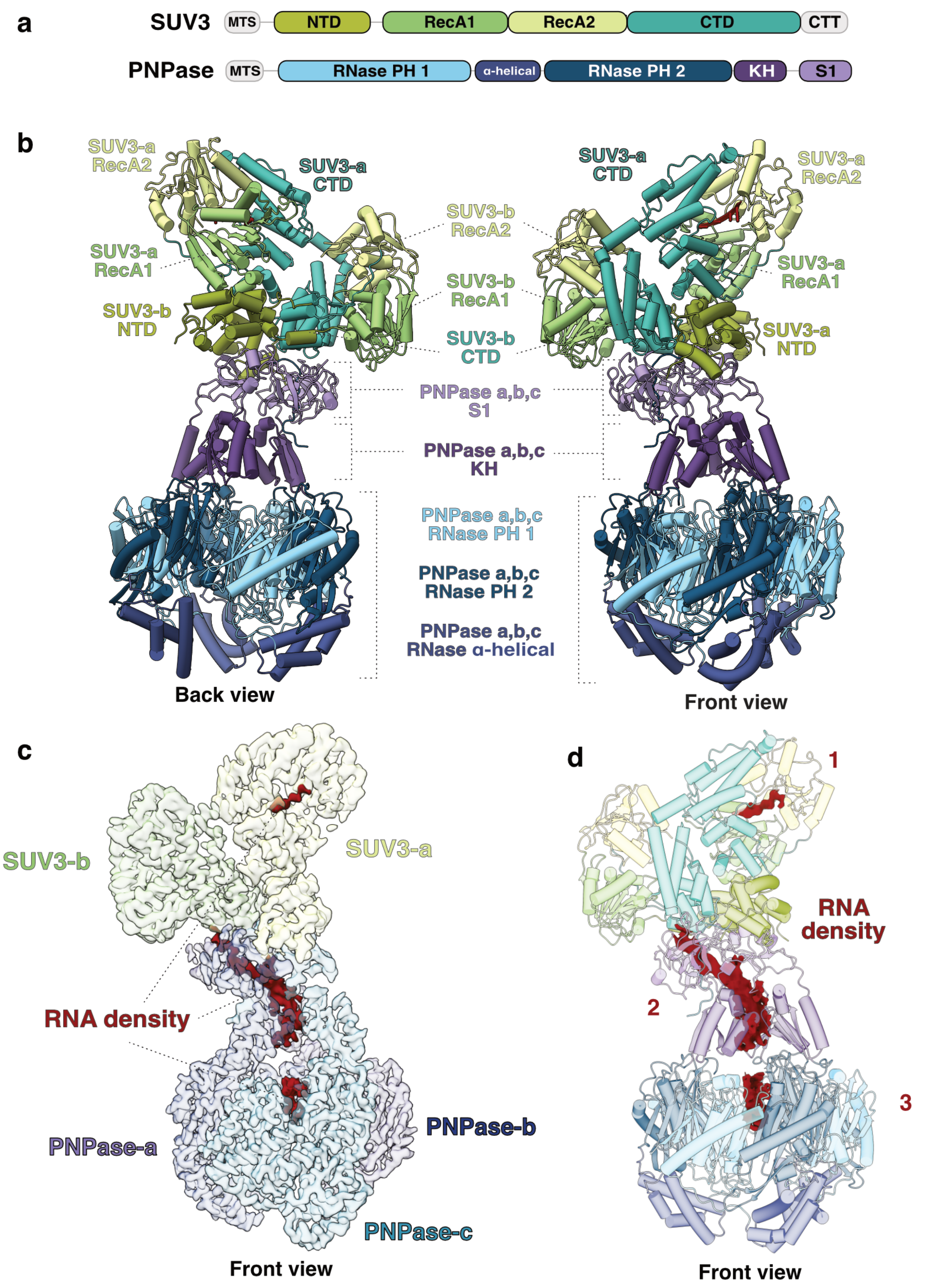


**Extended Data Figure 3 - Substrate-locked state of the SUV3–PNPase degradosome complex**

**a)** Schematic depiction of the primary structures and domain organization of SUV3 and PNPase.

**b)** Cartoon representation of the degradosome, with α-helices depicted as cylinders. SUV3 and PNPase are colored according to their domains, as indicated.

**c)** Cryo-EM map (Map D) showing RNA density within the degradosome. Regions corresponding to RNA are colored in red.

**d)** Cartoon representation of the degradosome, colored as in panels C–D, with α-helices depicted as cylinders. The cryo-EM map (Map D) regions corresponding to RNA are shown in red.

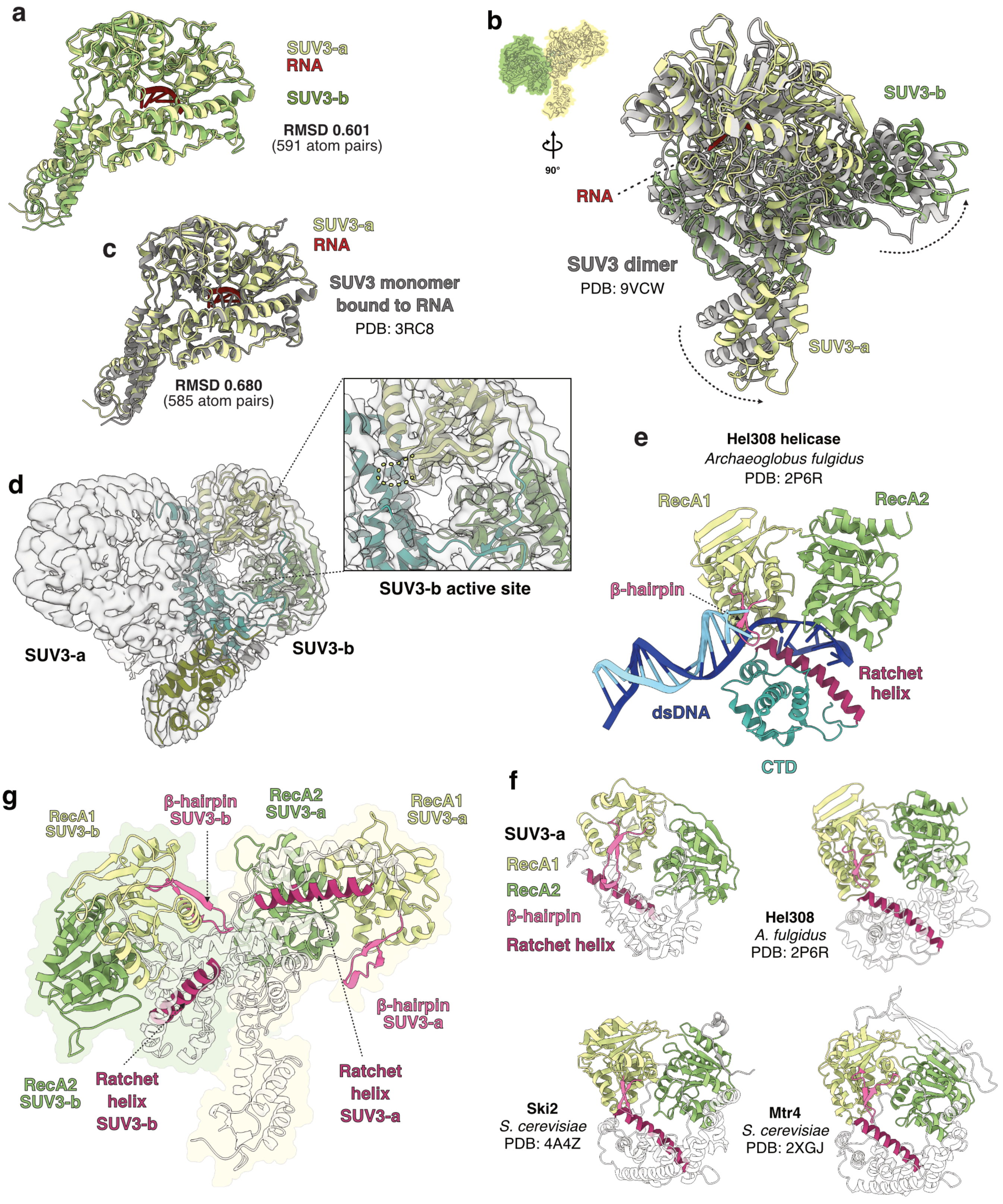


**Extended Data Figure 4 - Dimeric architecture and conserved features of SUV3 helicase**

Figure captions follow on the next page.

**Extended Data Figure 4 - Dimeric architecture and conserved features of SUV3 helicase**

**a)** Conserved architecture of SUV3 monomers within the degradosome-bound dimer. Structural alignment of SUV3-awith SUV3-b The RMSD value was calculated using SUV3-a as the reference structure (MatchMaker, ChimeraX [2]).

**b)** Structural comparison of the the SUV3 dimer bound to RNA within the human mitochondrial degradosome with the free SUV3 dimer bound to ssRNA (PDB: 9VCW [3]) and (colored by subunit). Dashed arrows illustrate conformational changes in the N-terminal regions of SUV3-a and SUV3-b relative to the helicase core.

**c)** Structural comparison of SUV3-a bound to RNA within the mitochondrial degradosome with the free SUV3 monomer bound to ssRNA (PDB: 3RC8 [4])

**d)** Focused cryo-EM reconstruction of the SUV3 dimer (Map C). The SUV3-b protomer is shown as a cartoon model. No additional density corresponding to bound RNA is observed within the active site of SUV3-b.

**e)** Model of the archaeal Hel308 helicase bound to a DNA duplex–single-stranded junction (PDB: 2P6R [5]). The RecA domains and C-terminal domain (CTD) are colored as in Figure 2A, while the ratchet helix and β-hairpin are shown in shades of pink. Parts of Hel308 CTD were omitted for clarity.

**f)** Structural comparison of the translocation module of SUV3-a with those of the SF2 helicases Hel308 (PDB: 2P6R [5]), Ski2 (PDB: 4A4Z [6]), and Mtr4 (PDB: 2XGJ [7]), highlighting conserved features of the nucleic acid translocation machinery.

**g)** Tandem arrangement of the translocation modules of SUV3-a and SUV3-b within the dimer. RNA in SUV3-a was omitted for clarity.

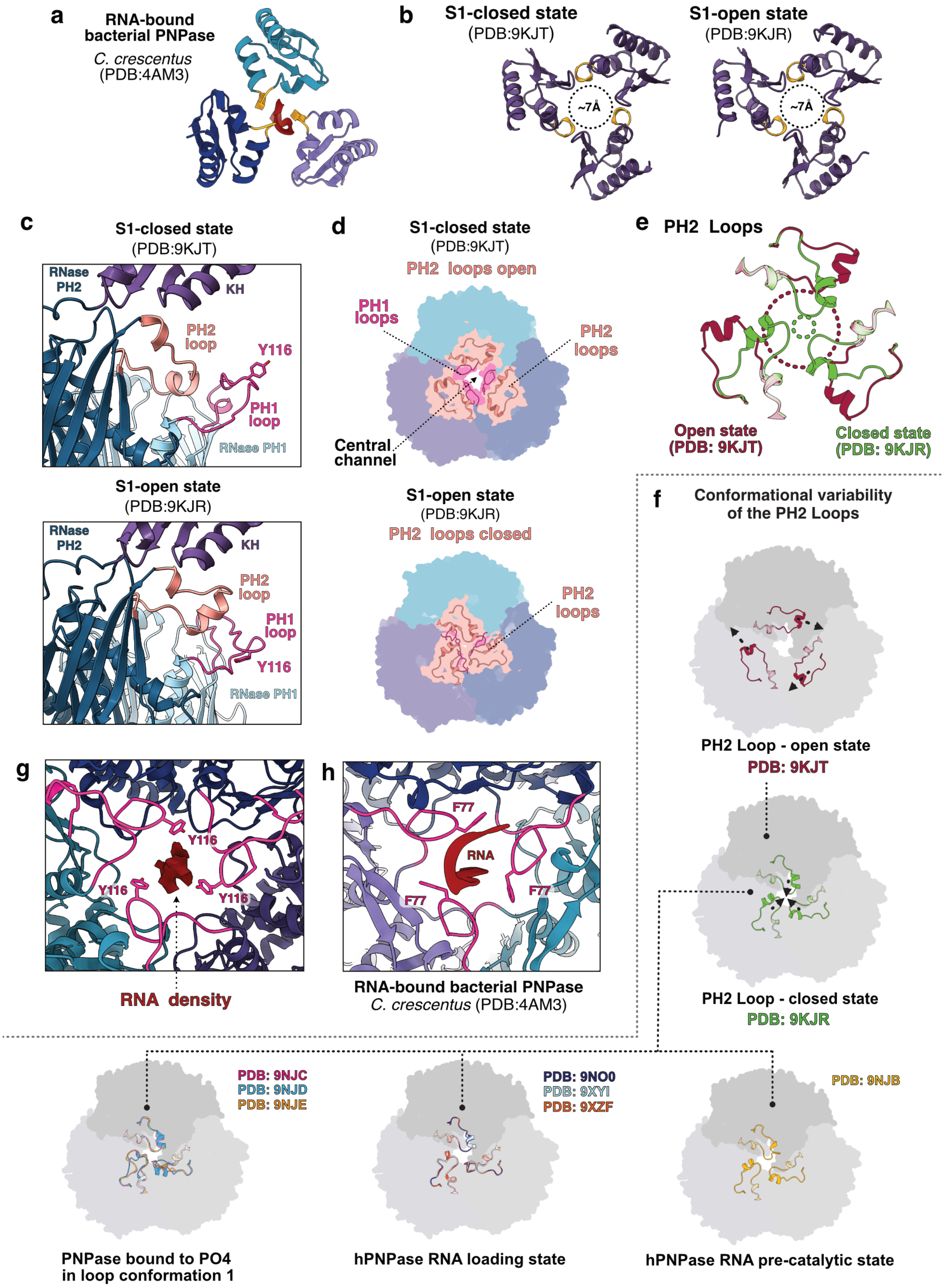


**Extended Data Figure 5 – Structural details of PNPase in the degradosome**

Figure captions follow on the next page.

**Extended Data Figure 5 – Structural details of PNPase**

**a)** KH domains of *C. crescentus* PNPase (PDB:4AM3 [9]) shown in the clamped conformation around an incoming ssRNA molecule.

**b)** Comparison of KH domain geometry in free PNPase in S1-closed state (PDB: 9KJT) and PNPase in S1-open state (PDB: 9KJR). Rearrangements of the KH domains in the degradosome (Figure 3G) constrict the central channel. The view is rotated relative to panel B.

**c)** Relative positions of the PH2 loop and PH1 loop in one PNPase protomer of the free PNPase on the open state (PDB: 9KJR [8]) and closed states of the S1 domains and (9KJT [8]). In degradosome PNPase (Figure 3H), the PH2 loop is disordered.

**d)** Relative positions of the PH2 loop in the open and closed conformations of PNPase (9KJT and 9KJR [8]). In the open conformation, the PH2 loop exposes the central RNA channel (9KJT [8]); in the closed conformation, it constricts the channel (9KJR [8]).

**e)** Superposition of the PH2 loops in the open and closed conformations of PNPase (PDB: 9KJT and 9KJR [8]). Dashed lines indicate the diameter of the central channel in each configuration.

**f)** Structural variability of the closed conformation of PH loop 2 in phosphate-bound PNPase (PDB: 9NJC, 9NJD, and 9NJE [10]), RNA-loading state (PDB: 9NO0, 9XYI, and 9XZF [10]), and the pre-catalytic state (PDB: 9NJB [10]).

**g)** Side-by-side comparison of RNA coordination in *C. crescentus* PNPase (PDB:4AM3 [9]) and hPNPase. In *C. crescentus* PNPase, RNA is coordinated by F77 of the FLRR loop, whereas in hPNPase, RNA density (Map D) is observed in the vicinity of Y116 of the YLRR motif within the PH1 loop.

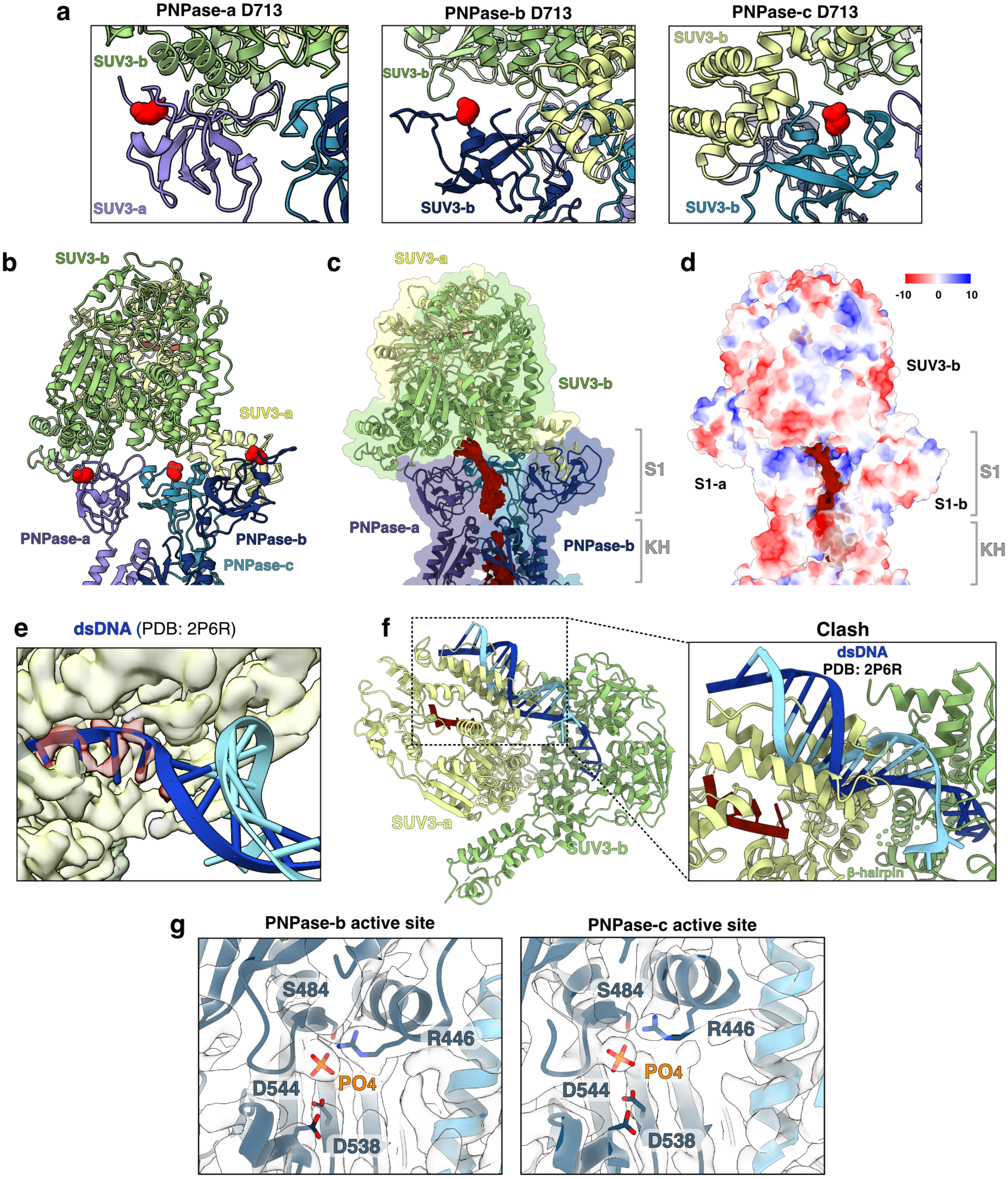


**Extended Data Figure 6 - Interactions of the human helicase SUV3 with exonucleases and model of RNA interactions in the degradosome complex**

Figure captions follow on the next page.

**Extended Data Figure 6 – Pathogenic mutations impairing degradosome formation and model of RNA interactions within the degradosome complex**

**a)** Close-up view of D713 (shown as red spheres) in each PNPase protomer.

**b)** Side view of the degradosome complex shown as a cartoon representation, highlighting the location of D713 (red spheres) in all three PNPase protomers. In each protomer, D713 is positioned in close proximity to the SUV3-binding interface**.**

**c)** Side view of the degradosome complex shown as molecular surfaces, colored according to electrostatic potential (red, negative; blue, positive). The dark red density (Map D) corresponds to RNA accommodated within a positively charged cleft of the complex.

**d)** Same view as in panel A, with the complex shown as cartoon. The dark red region represents RNA density in a low-threshold cryo-EM map (Map D).

**e)** Superposition of the archaeal helicase Hel308 bound to dsDNA (PDB: 2P6R [5]) onto SUV3-a. The single-stranded 3′ overhang of the nucleic acid aligns precisely with the RNA density observed at the SUV3-a active site (Map D). SUV3-a and Hel308 are omitted for clarity.

**f)** Superimposition of the archaeal helicase Hel308 (PDB: 2P6R [5]) bound to dsDNA onto SUV3-b. The modeled nucleic acid would sterically clash with SUV3-a, as shown in the zoomed-in view. The unwound DNA strand is colored dark blue and the complementary strand light blue.

**g)** Detailed view of the active sites of PNPase b and c, highlighting the catalytic residues. The phosphate ion was positioned based on comparison with the apo structure of *C. crescentus* PNPase (PDB: 4AIM [61]).

**Extended Data Table 1 - RNA oligonucleotides used in this study -** RNA oligonucleotides and annealed products used for RNA degradation assays and Cryo-EM sample preparation.

<table>
<tr><th colspan="3">RNA probes used for degradation assays and cryo-EM analysis</th></tr>
<tr><th>Oligo ID</th><th>Oligo sequence</th><th>Annealed oligos</th></tr>
<tr><td>ssRNA_01</td><td>5' /56-FAM/AAGUGGUCGAUGUAGUAGUGUGAGUGAGUGUGCAGAUCAGAGACUUG 3'</td><td></td></tr>
<tr><td>ssRNA_02</td><td>5' CACACUACUACAUCGACCACUU 3'</td><td></td></tr>
<tr><td>ssRNA_03</td><td>5' /56-FAM/GGCCGCGGAAGUGGUCGAUGUAGUAGUGUGAGUGAGUGUGCCCCAGAUCAGUGGUAGUGU 3'</td><td></td></tr>
<tr><td>ssRNA_04</td><td>5' /56-FAM/GGCCGCGGAAG*U*G*G*U*CGAUGUAGUAGUGUGAGUGAGUGUGCCCCAGAUCAGUGGUAGUGU 3'</td><td></td></tr>
<tr><td>ssRNA_05</td><td>5' UUCCGCGGCC 3'</td><td></td></tr>
<tr><td>dsRNA_01</td><td>5' /56-FAM/AAGUGGUCGAUGUAGUAGUGUGAGUGAGUGUGCAGAUCAGAGACUUG 3'<br>||||||||||||||||||||||<br>3' UUCACCAGCUACAUCAUCACAC 5'</td><td>ssRNA_01<br>+<br>ssRNA_02</td></tr>
<tr><td>dsRNA_02_WT</td><td>5' /56-FAM/GGCCGCGGAAGUGGUCGAUGUAGUAGUGUGAGUGAGUGUGCCCCAGAUCAGUGGUAGUGU 3'<br>||||||||||<br>3' CCGGCGCCUU 5'</td><td>ssRNA_03<br>+<br>ssRNA_05</td></tr>
<tr><td>dsRNA_02_PT</td><td>5' /56-FAM/GGCCGCGGAAG*U*G*G*UCGAUGUAGUAGUGUGAGUGAGUGUGCCCCAGAUCAGUGGUAGUGU 3'<br>||||||||||<br>3' CCGGCGCCUU 5'</td><td>ssRNA_04<br>+<br>ssRNA_05</td></tr>
</table>

**Extended Data Table 2 – Cryo-EM data collection, refinement and validation statistics**

| **Data collection and processing** | |
|---|---|
| Magnification | 105,000 x |
| Voltage (kV) | 300 |
| Electron exposure (e–/Å$^2$) | 50 |
| Defocus range (μm) | 0.5 – 2.5 |
| Pixel size (Å) | 0.834 |
| Symmetry imposed | C1 |
| Initial particle images (no.) | 16,653,675 |

| | **Map A Consensus Map** | **Map B (PNPase)** | **Map C (SUV3)** | **Map C Composite Map** |
|---|---|---|---|---|
| **EMDB ID** | **EMD-58691** | **EMD-58689** | **EMD-58690** | **EMD-58688** |
| Final particle images (no.) | 209.892 | 209.892 | 209.892 | - |
| Map resolution (Å) | 3.3 | 3.2 | 3.6 | - |
| FSC threshold | 0.143 | 0.143 | 0.143 | - |
| Map sharpening *B* factor (Å$^2$) | -131.26 | -115.05 | -136.50 | - |
| **Refinement** | **PDB: 31WW** | | | |
| Model resolution (Å) | 3.4 | | | |
| FSC threshold | 0.143 | | | |
| Model composition | | | | |
| Non-hydrogen atoms | 25905 | | | |
| Protein residues | 3303 | | | |
| Ligands | | | | |
| *B* factors (Å$^2$) | | | | |
| Protein | 115.72 | | | |
| Ligand | | | | |
| R.m.s. deviations | | | | |
| Bond <lengths (Å) | 0.005 | | | |
| Bond angles (°) | 0.723 | | | |
| Validation | | | | |
| MolProbity score | 1.54 | | | |
| Clashscore | 6.29 | | | |
| Poor rotamers (%) | 0.98 | | | |
| Ramachandran plot | | | | |
| Favored (%) | 96.80 | | | |
| Allowed (%) | 3.11 | | | |
| Disallowed (%) | 0.09 | | | |

**Extended Data Table 3 - DNA oligonucleotides used in this study -** DNA oligonucleotides used for cloning and generation of SUV3 and PNPase constructs.

| **Oligonucleotides used for cloning** | | |
|---|---|---|
| **Oligo ID** | **Oligo sequence (5' - 3')** | **Cloning method** |
| pETSUMO_Δ46SUV3_Forward | 5' /5Phos/ACCGCCTCCTCCTCTGCC 3' | Restriction cloning |
| pETSUMO_Δ46SUV3_**NotI**_STOP_Reverse | 5' TGACT**GCGGCCGC**TTAGTCCGAATCAGGTTCCTTCTTC 3' | Restriction cloning |
| 14A_ **LIC-v2F**_Δ39PNPase_ Forward | 5' **TTTAAGAAGGAGATATAGATC**ATGAGTAGCGCAGGGTCTCGA 3' | Ligation independent |
| 14A_Δ39PNPase-His_ **LIC-v2R**_Reverse | 5' **TTATGGAGTTGGGATCTTATTA**GTGGTGGTGGTGGTGGTGCTCGAGTGC 3' | Ligation independent |
| 438A_**LIC-vBacF**_Met_Δ23PNPase_Forward | 5' **TACTTCCAATCCAATCG**ATGCCACGGCGGGATCGGGCAC 3' | Ligation independent |
| 438A_PNPase-His_**vLIC-v1rv**_Reverse | 5' **TTATCCACTTCCAATGTTATTA**GTGGTGGTGGTGGTGGTGCTCGAGTGC 3' | Ligation independent |

### Extended Data References


1. Tegunov, D., and Cramer, P. (2019). Real-time cryo-electron microscopy data preprocessing with Warp. Nat. Methods *16*, 1146–1152. https://doi.org/10.1038/s41592-019-0580-y.

2. Pettersen, E.F., Goddard, T.D., Huang, C.C., Meng, E.C., Couch, G.S., Croll, T.I., Morris, J.H., and Ferrin, T.E. (2021). UCSF ChimeraX: Structure visualization for researchers, educators, and developers. Protein Sci. *30*, 70–82. https://doi.org/10.1002/pro.3943.

3. Patra, M., Jain, M., Li, Y.-C., Chen, Y.-P., Golzarroshan, B., and Yuan, H.S. (2026). Asymmetric dimeric assembly of Suv3 helicase facilitates processive RNA unwinding. Nat. Commun. https://doi.org/10.1038/s41467-026-71901-2.

4. Jedrzejczak, R., Wang, J., Dauter, M., Szczesny, R.J., Stepien, P.P., and Dauter, Z. (2011). Human Suv3 protein reveals unique features among SF2 helicases. Acta Crystallogr. Sect. D *67*, 988–996. https://doi.org/10.1107/s0907444911040248.

5. Büttner, K., Nehring, S., and Hopfner, K.-P. (2007). Structural basis for DNA duplex separation by a superfamily-2 helicase. Nat. Struct. Mol. Biol. *14*, 647–652. https://doi.org/10.1038/nsmb1246.

6. Halbach, F., Rode, M., and Conti, E. (2012). The crystal structure of S. cerevisiae Ski2, a DExH helicase associated with the cytoplasmic functions of the exosome. RNA *18*, 124–134. https://doi.org/10.1261/rna.029553.111.

7. Weir, J.R., Bonneau, F., Hentschel, J., and Conti, E. (2010). Structural analysis reveals the characteristic features of Mtr4, a DExH helicase involved in nuclear RNA processing and surveillance. Proc. Natl. Acad. Sci. *107*, 12139–12144. https://doi.org/10.1073/pnas.1004953107.

8. Li, Y.-C., Wang, C.-H., Patra, M., Chen, Y.-P., Yang, W.-Z., and Yuan, H.S. (2025). Structural insights into human PNPase in health and disease. Nucleic Acids Res. *53*, gkaf119. https://doi.org/10.1093/nar/gkaf119.

9. Hardwick, S.W., Gubbey, T., Hug, I., Jenal, U., and Luisi, B.F. (2012). Crystal structure of Caulobacter crescentus polynucleotide phosphorylase reveals a mechanism of RNA substrate channelling and RNA degradosome assembly. Open Biol. *2*, 120028. https://doi.org/10.1098/rsob.120028.

10. Unseld, O., Das, H., and Hällberg, B.M. (2025). Loop-mediated regulation and base flipping drive RNA cleavage by human mitochondrial PNPase. Nucleic Acids Res. *53*, gkaf1296. https://doi.org/10.1093/nar/gkaf1296.

11. Razew, M., Warkocki, Z., Taube, M., Kolondra, A., Czarnocki-Cieciura, M., Nowak, E., Labedzka-Dmoch, K., Kawinska, A., Piatkowski, J., Golik, P., et al. (2018). Structural analysis of mtEXO mitochondrial RNA degradosome reveals tight coupling of nuclease and helicase components. Nat. Commun. *9*, 97. https://doi.org/10.1038/s41467-017-02570-5.

12. Weick, E.-M., Puno, M.R., Januszyk, K., Zinder, J.C., DiMattia, M.A., and Lima, C.D. (2018). Helicase-Dependent RNA Decay Illuminated by a Cryo-EM Structure of a Human Nuclear RNA Exosome-MTR4 Complex. Cell *173*, 1663-1677.e21. https://doi.org/10.1016/j.cell.2018.05.041.
